# KAI2-dependent signaling controls vegetative reproduction in *Marchantia polymorpha* through activation of LOG-mediated cytokinin synthesis

**DOI:** 10.1101/2024.09.02.610783

**Authors:** Aino Komatsu, Mizuki Fujibayashi, Kazato Kumagai, Hidemasa Suzuki, Yuki Hata, Yumiko Takebayashi, Mikiko Kojima, Hitoshi Sakakibara, Junko Kyozuka

## Abstract

*Marchantia polymorpha* reproduces vegetatively (asexually) by producing propagules known as gemmae within gemma cups and sexually through spores. We previously reported that KARRIKIN INSENSITIVE 2 (KAI2)-dependent signaling promotes gemma cup and gemma formation. KAI2A perceives unidentified endogenous ligand(s), tentatively referred to as KAI2 ligands (KL). Perception of KL by KAI2 triggers MAX2-dependent proteolysis of MpSMXL. In this study, we identified genes working downstream of KAI2-dependent signaling in *M. polymorpha*. We found that KAI2-dependent signaling positively controls the expression of Mp*LONLEY GUY* (Mp*LOG*), encoding a cytokinin biosynthesis enzyme. Disruption of the Mp*LOG* function decreased endogenous cytokinin levels and caused defects similar to KAI2-dependent signaling mutants. Moreover, supplying exogenous cytokinins rescued the defects of Mp*log* and KAI2-dependent signaling mutants, implying that cytokinins work downstream of KAI2-dependent signaling. Activation of Mp*LOG* by KAI2-dependent signaling occurs in a highly cell-type-specific manner, leading to cell-specific induction of *GEMMA CUP-ASSOCIATED MYB1* (*GCAM1*), the master regulator of vegetative reproduction of *M. polymorpha*. We propose a genetic cascade, starting from KAI2-dependent signaling, that promotes vegetative reproduction through the induction of Mp*LOG* and *GCAM1*. The interaction between KAI2-dependent signaling and cytokinin in *M. polymorpha* provides a novel insight into the function and evolution of KAI2-dependent signaling.

## Introduction

A wide range of plants uses vegetative or asexual reproduction for propagation as their predominant mechanisms of reproduction^1^. In vegetative reproduction, new plants are produced via propagules, such as adventitious shoots, bulbs, tubers, and rhizome buds^2^. In addition, new individuals can be formed through fragmentation and clonal growth of the original plant^3, 4^. Vegetative reproduction is less affected by seasonal changes than sexual reproduction, which is often influenced by seasonal factors such as light. Consequently, vegetative reproduction allows efficient propagation, potentially yielding a huge number of individuals. At the same time, vegetative reproduction has a clear disadvantage in that it limits the potential for increasing genetic diversity. Plants regulate the degree of vegetative reproduction in response to environmental conditions to optimize proliferation, and the molecular mechanisms underlying this regulation are mostly unknown. In bryophytes, new individuals are generated non-sexually from propagules and caducous organs of diverse morphology^2, 5, 6^. In addition, the high regeneration capacity of bryophytes enables the generation of new individuals from fragments or parts of gametophytes, such as leaves, shoots, and thalli. *Marchantia polymorpha* proliferates both through sexual and vegetative reproduction^3^. For vegetative reproduction, propagules called gemmae are formed on the base of the gemma cup serving as the primary source for extensive propagation^7^.

Plant hormones are key regulators of almost all aspects of plant growth and development as well as responses to environmental conditions^8, 9^. Fine-tuning of synthesis and signaling is pivotal for controlling plant hormone function. The hormonal actions are intricately interconnected, and the crosstalks among plant hormones are crucial to maximize their effects^10^. We previously reported that gemma cup and gemma formation are positively controlled by a KARRIKIN INSENSITIVE 2 (KAI2)-dependent signaling pathway in *M. polymorpha*^11, 12^. KAI2 was identified as the receptor for karrikins (KARs), molecules derived from smoke from burning vegetation^13, 14^. Subsequent genetic analyses have suggested that KAI2 most likely perceives endogenous ligand(s), currently unidentified and provisionally termed KAI2 ligands (KL)^15^. The KAR/KL signal is transduced through the MORE AXILLARY GROWTH 2 (MAX2)-mediated proteolysis-dependent signaling pathway. In this pathway, ligand perception by KAI2 triggers the interaction of MAX2 with suppressor proteins, resulting in the degradation of these suppressor proteins by the 26S proteasome pathway^14, 16, 17, 18, 19^. In *Arabidopsis*, SUPPRESSOR OF MAX2 1 (SMAX1) and SMAX1 LIKE2 (SMXL2), encoding proteins with weak homology to ClpB-type chaperonins, work as the suppressors of KAI2-dependent signaling^16, 20, 21^. SMXL6, SMXL7, and SMXL8, paralogs of SMAX1, function as suppressors of the strigolactone signaling pathway^16, 21,22^. All SMXL proteins contain an ethylene-responsive element binding factor-associated amphiphilic repressor (EAR) motif that interacts with transcriptional corepressors, including TOPLESS (TPL). Consequently, SMXLs are presumed to work as transcription repressors, engaging with transcription factors and transcriptional corepressors^23, 24^. In addition, SMXL6 works as a transcription factor that suppresses SMXL 6, 7, and 8 transcription through direct binding to their promoters^23^. Downstream targets of transcriptional repression mediated by SMXL corepressors, such as *BRANCHIED1*, *TCP1*, and *PRODUCTION OF ANTHOCYANIN PIGMENT1*, have been identified^23, 24^. A few transcription factors that directly interact with SMXLs have been identified, however, the comprehensive understanding of gene expression regulation by SMXLs is yet to be revealed ^25, 26^.

The KAI2-dependent signaling pathway in seed plants controls diverse aspects of growth and development, including seed germination, seedling growth, leaf development, root growth, root hair development, stress tolerance, and arbuscular mycorrhizal symbiosis^20, 21, 27, 28, 29, 30, 31^. In addition, crosstalks of KAI2-dependent signaling with other hormones allow for an elaborate fine-tuning of growth and development and responses to the ever-changing environment^32^. In *Arabidopsis* and *Lotus japonica*, KAI2-dependent signaling controls root and root hair growth in response to low P conditions through modulation of ethylene biosynthesis and signaling^33, 34^. This leads to the modulation of spatial accumulation of PIN2 and AUX1, leading to changes in auxin accumulation patterns. Shade avoidance in *Arabidopsis* is also mediated by KAR/KL-auxin crosstalk^28, 35^. Interaction of KAI2-dependent signaling and ABA contributes to the control of drought tolerance and seed germination^36^. KAI2-dependent signaling interacts with strigolactone biosynthesis, which leads to the regulation of arbuscular mycorrhizal (AM) fungi colonization in rice^31^.

All the core components of KAI2-dependent signaling exist in bryophytes^37, 38^. *M. polymorpha* contains two KAI2 paralogues (Mp*KAI2A* and Mp*KAI2B*), one *MAX2* gene (Mp*MAX2*), and one *SMXL* (Mp*SMXL*). We previously reported that Mp*KAI2A*, Mp*MAX2*, and Mp*SMXL* work in a single signaling pathway, which depends on the degradation of MpSMXL^12^. Although MpKAI2B shows extensive sequence similarity to MpKAI2A, it is unlikely that MpKAI2B works as the KAR/KL signaling receptor. The conservation of the KAI2-dependent signaling pathway in bryophytes was demonstrated in the moss *Physcomitrium patens*, and the liverworts *M. polymorpha* and *Marchantia paleacea*, indicating that KAI2-dependent signaling was established in the common ancestor of land plants^12, 39, 40^. Later, in the common ancestor of seed plants, the *KAI2* gene was duplicated, which led to the evolution of D14, a strigolactone receptor^14, 41^. Strigolactone signals are transduced via a parallel signaling pathway that shares MAX2 with KAI2-dependent signaling^16, 17, 18, 42^. Other than the occurrence of *D14*, the increase in the number of *KAI2* paralogs occurred independently in some lineages, such as the root parasitic plants and *P. patens*^41, 42, 43, 44, 45^.

We are interested in uncovering the mechanisms by which KAI2-dependent signaling controls vegetative reproduction in *M. polymorpha*. In this study we identified genes working downstream of KAI2-dependent signaling in *M. polymorpha*. Our analysis indicates that KAI2-dependent signaling positively controls the expression of Mp*LOG*, encoding a cytokinin biosynthesis enzyme^46^. We present evidence that cytokinin works downstream of KAI2-dependent signaling. We also demonstrate that activation of Mp*LOG* by KAI2-dependent signaling occurs in a highly cell-type-specific manner, leading to cell-specific induction of *GEMMA CUP-ASSOCIATED MYB1* (*GCAM1*), the master regulator of vegetative reproduction^11, 47^. The cross-regulation between KAI2-dependent signaling and cytokinin in *M. polymorpha* provides a novel insight into the function and evolution of KAI2-dependent signaling.

## Results

### MpLOG works downstream of KAI2-dependent signaling

We first aimed at identifying genes working downstream of KAI2-dependent signaling by analyzing transcriptome changes upon the induction of MpMAX2-dependent degradation of MpSMXL. For this purpose, we took advantage of the fact that *M. polymorpha* does not respond to strigolactones due to the absence of D14, the cognate receptor of strigolactones. We previously reported that *M. polymorpha* and its close relative *M. paleacea* respond to (‒)-GR24, but not to (+)-GR24, synthetic compounds mimicking KL and strigolactones, respectively. However the introduction of *AtD14* enabled *M. polymorpha* and *M. paleacea* to respond to both (+)-GR24 and (‒)-GR24 with the signals transmitted through MpMAX2-dependent degradation of MpSMXL^39^. In *35S:AtD14* transgenic lines of *M. polymorpha* (Mp*AtD14ox*), MpSMXL degradation occurs upon perception of endogenous KL and applied (‒)-GR24 by MpKAI2A, and perception of applied (+)-GR24 by the introduced AtD14. In addition, since *AtD14* is driven by the strong and constitutive 35S promoter, and AtD14-dependent signaling occurs only after the addition of *rac*-GR24 containing both (+)-GR24 and (‒)-GR24, we anticipate that gene expression changes would be amplified in the Mp*AtD14ox* line, making it suitable for RNAseq analysis. We applied *rac*-GR24 to the Mp*AtD14ox* line and compared changes in transcriptomes 3 and 6 hours after application (Fig. 1a). We identified 283 and 402 genes that were up-regulated and 182 and 278 genes that were down-regulated, 3 and 6 hours respectively, after the application of *rac*-GR24. Among the differentially expressed genes (DEGs), 76 genes were up-regulated and 43 genes were down-regulated, at both 3 and 6 hours (Fig. 1b and Supplementary Fig. 1a).

**Figure 1.**
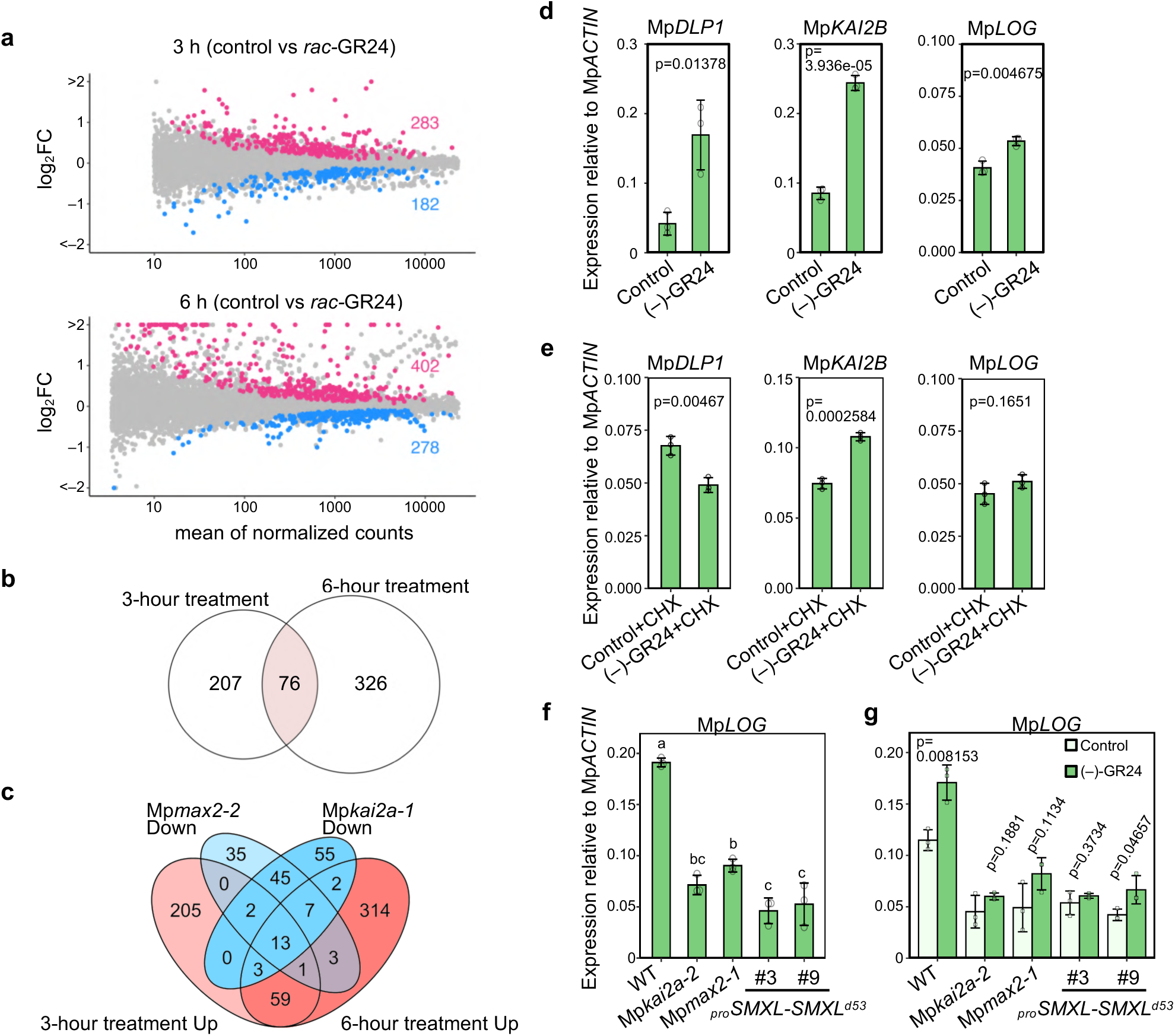
MpLOG works downstream of KAI2-dependent signaling. **a** MA plots showing the differential gene expression in 1 µM *rac-*GR24-treated Mp*AtD14ox* plants compared to mock-treated Mp*AtD14ox* plants, 3 hours (top) and 6 hours (bottom) after the start of treatment. **b** Venn diagram showing the number of up-regulated genes after 3 and 6 hours of 1 µM *rac-*GR24 treatment in Mp*AtD14ox* plants. **c** Venn diagram showing the number of up-regulated genes after 1 µM *rac-*GR24 treatment in Mp*AtD14ox* plants and the down-regulated genes in Mp*max2-2* and Mp*kai2a-1* mutants. **d** Induction of three up-regulated genes (Mp*DLP1*, Mp*KAI2B*, and Mp*LOG*) in WT by 1 µM (–)-GR24 treatment for 6 hours examined by quantitative PCR (qPCR) analysis. **e** Induction of Mp*DLP1*, Mp*KAI2B*, and Mp*LOG* expression by 1 µM (–)-GR24 treatment for 6 hours in the presence of 1 µM Cycloheximide (CHX) examined by qPCR analysis in WT. **f** Expression level of Mp*LOG* in WT, Mp*kai2a-2*, Mp*max2-1*, *_pro_SMXL-SMXL^d53^*#3 and *_pro_SMXL-SMXL^d53^*#9. **g** Induction of Mp*LOG* expression by 1 µM (–)-GR24 treatment for 6 hours in WT, Mp*kai2a-2*, Mp*max2-1*, *_pro_SMXL-SMXL^d53^*#3, and *_pro_SMXL-SMXL^d53^*#9 examined by qPCR analysis. Expression levels relative to *ACTIN* (Mp*ACTIN*) are shown (**d-g)**. Bars represent means ± SD (n=3). The p values were calculated by using the Student’s t-tests (**d, e, g)**. The Tukey’s HSD test was used for multiple comparisons in **f** and statistical differences (p<0.05) are indicated by different letters.

GO analysis identified that synthesis and metabolism of small compounds were enriched in DEGs up-regulated after 3 hours. In contrast, GO terms related to transcription and replication were enriched in DEGs down-regulated after 3 hours. On the other hand, GO terms related to growth and differentiation were enriched in DEGs down-regulated after 6 hours, while no specific GO term was identified in DEGs up-regulated after 6 hours (Supplementary Fig. 1c, d).

In our previous study, we identified genes whose expression changed in Mp*kai2a* and Mp*max2*, mutants lacking KAI2-dependent signaling^12^. However, due to advancements in gene annotation of *M. polymorpha* progressed, we re-analyzed the RNAseq data and identified additional genes as DEGs. We integrated the data from two comparisons: WT versus mutants and non-treated Mp*AtD14ox* versus *rac*-GR24-treated Mp*AtD14ox*. We found that the expression levels of 13 genes was increased by *rac*-GR24 treatment after 3 and 6 hours and decreased in the Mp*kai2a* and Mp*max2* mutants (Fig. 1c, Supplementary Fig. 1b), suggesting that these genes are likely positively regulated by KAI2-dependent signaling.

First, we confirmed that the expression of 12 out of the 13 DEGs is enhanced in WT plants after application of (‒)-GR24 (Fig. 1d, Supplementary Fig. 2). We picked three genes, Mp*DIENELACTONE HYDROLASE LIKE PROTEIN1* (Mp*DLP1)*, Mp*KAI2B*, and Mp*LOG*, from the 13 DEGs for further analysis (Supplementary Table 1). The expression of genes controlled by transcription factors that directly interact with MpSMXL is expected to change without the need for a translation step. Indeed, Mp*KAI2B*, but not the other two genes, was activated by (‒)-GR24 in the presence of cycloheximide (CHX), a translation repressor, suggesting that Mp*KAI2B* might be the direct target of KAI2-dependent signaling (Fig. 1e).

In this study, we focused on Mp*LOG*, an ortholog of *LONELY GUY* encoding the final enzyme in the active cytokinin biosynthesis pathway^46, 48, 49^. Mp*LOG*, a single-copy gene in *M. polymorpha*, shows high sequence similarity to *LOG* genes in other plant species across the entire coding sequence. We confirmed that Mp*LOG* expression is reduced in the Mp*kai2a*, Mp*max2* mutants, as well as in transgenic lines overexpressing Mp*SMXL^d53^*, which encodes a degradation-resistant version of the MpSMXL suppressor protein (Fig. 1f)^12^. Moreover, induction of Mp*LOG* expression by (‒)-GR24 observed in WT was abolished in lines with the defective KAI2-dependent signaling (Fig. 1g). These results suggest that KAI2-dependent signaling positively controls Mp*LOG* expression.

### MpLOG catalyzes the final steps of cytokinin biosynthesis in *M. polymorpha*

The cytokinin biosynthesis pathways in plants have been elucidated through studies in angiosperms (Fig. 2a)^48, 49^. *LOG* encodes a cytokinin riboside 5′-monophosphate phosphoribohydrolase that releases cytokinin nucleobase and ribose 5′-monophosphate from iPRMP, tZRMP, DZRMP, and cZRMP, leading to the production of N^6^-isopentenyl adenine (iP), *trans*-zeatin (tZ), dihydro-zeatin (DZ), and *cis*-zeatin (cZ), all active cytokinins^46^. Phylogenetic studies demonstrated that *M. polymorpha* contains a single *LOG* gene (Supplementary Fig. 3) while it does not contain genes for canonical adenosine phosphate-isopentenyltransferase (IPT) and CYP735A. It was shown that DZ content is below the detectable level in *M. polymorpha*^50^.

**Figure 2.**
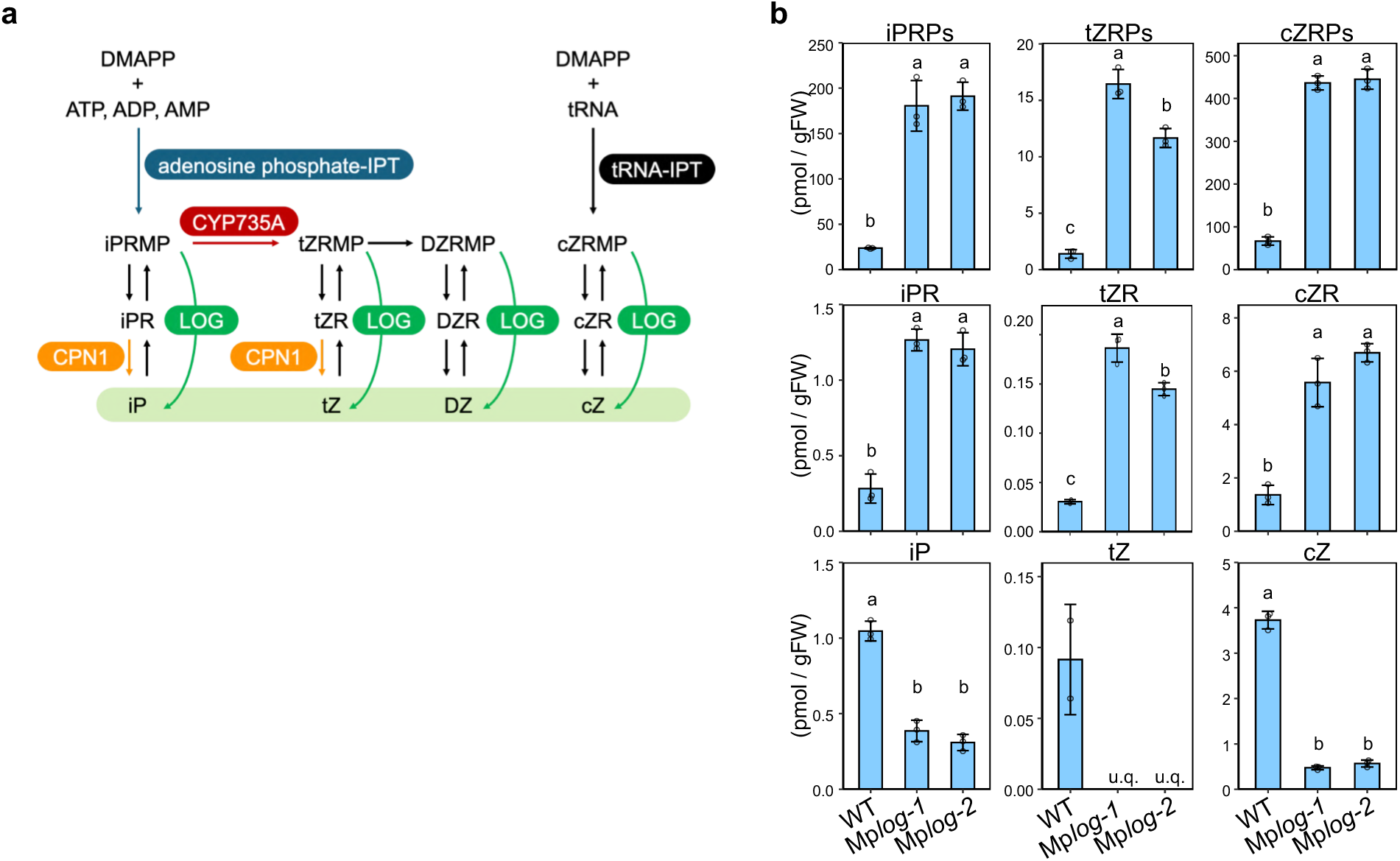
MpLOG catalyzes the final steps of cytokinin biosynthesis in *M. polymorpha*. **a** Cytokinin biosynthesis pathway in angiosperms. The addition of an isoprenoid side chain to the N^6^-amino group of an adenine nucleotide by isopentenyltransferases (IPTs) is the rate-limiting step in cytokinin synthesis^33^. Two different forms of IPTs, adenosine phosphate-IPTs and tRNA-IPTs, are known in plants. Adenosine phosphate-IPTs use ATP, ADP, or AMP as substrates for prenylation. Subsequently, with the function of CYP735A and an unidentified zeatin reductase, iP riboside 5′-monophosphate (iPRMP), *trans-*zeatin 5′-monophosphate (tZRMP), and dihydro zeatin 5′-monophosphate (DZRMP) are synthesized. tRNA-IPTs prenylate UNN-decoding tRNAs at an adenine (A_37_), and the release of cytokinin nucleotides through tRNA degradation produces *cis-*zeatin 5′-monophosphate (cZRMP). LOG encodes a cytokinin riboside 5′-monophosphate phosphoribohydrolase that releases cytokinin nucleobase and ribose 5′-monophosphate from iPRMP, tZRMP, DZRMP, and cZRMP, leading to the production of N^6^-isopentenyl adenine (iP), *trans*-zeatin (tZ), dihydro-zeatin (DZ), and *cis*-zeatin (cZ), active cytokinins. Cytokinin/purine riboside nucleosidase1 (CPN1) catalyzes the deribosylation of cytokinin riboside precursors and other purine ribosides. **b** Quantification of cytokinins in WT, Mp*log-1*, and Mp*log-2*. Bars represent means ± SD (n=3). The Tukey’s HSD test was used for multiple comparisons and statistical differences (p<0.05) are indicated by different letters. u.q. indicates under quantification limits. iPRPs, tZRPs, and cZRPs; ribotide precursors, iPR, tZR, and cZR; riboside precursors.

To examine if Mp*LOG* works in the biosynthesis of cytokinins in *M. polymorpha*, we generated loss-of-function mutants of Mp*LOG* using the CRISPR/Cas9 genome editing system with two different gRNAs (Supplementary Fig. 4a) and subsequently measured the cytokinin contents of the resulting plants. We found that cZ was the most abundant cytokinin in WT *M. polymorpha* plants. iP and tZ were also detected (Fig. 2b). The content of tZ was reduced to undetectable levels in the Mp*log* mutants. The levels of iP and cZ were also significantly decreased; however, low levels of iP and cZ were detected in the Mp*log* mutants. In contrast to the decrease in active cytokinins, their ribotide precursors, iPRP, tZRP, and cZRP, were dramatically increased. The riboside precursors, iPR, tZR, and cZR, were also increased, but the amount was two orders of magnitude less than the ribotide precursors. These results support the notion that MpLOG catalyzes the final two steps in cytokinin biosynthesis. In addition, the presence of active cytokinins even in Mp*log* mutants strongly suggests the existence of a MpLOG-independent cytokinin synthesis pathway.

### Cytokinins synthesized by Mp*LOG* control gemma cup and gemma formation, as well as thallus growth

We previously reported that KAI2-dependent signaling controls gemma cup and gemma formation, flat thallus growth, and gemma growth arrest in the dark ^11, 12, 39^. We analyzed the phenotype of Mp*log* loss-of-function mutants to investigate whether Mp*LOG* is involved in the regulation of these traits under KAI2-dependent signaling. We reported that gemma cup formation in KAI2-dependent signaling mutants is affected by culture conditions. Specifically, gemma cup formation appears normal on media, whereas it is significantly suppressed when plants are grown on vermiculite ^11, 12^. Therefore, we analyzed gemma cup formation in Mp*log* mutant plants grown on vermiculite. Gemma cup formation was severely suppressed in these plants (Fig. 3a, b), and these defects were rescued by adding 6-benzylaminopurine (BA), a synthetic cytokinin. On the other hand, when Mp*log* mutant plants were grown on medium, gemma cups and gemmae were formed, but the number of gemmae formed per cup was much lower than that in the WT, similar to what is observed in KAI2 signaling mutants (Fig. 3c, d)^11, 12^. These results suggest that MpLOG-derived cytokinin positively controls gemma cup and gemma formation, resembling the function of genes in the KAI2-dependent signaling pathway. Gemmae of Mp*log* loss-of-function mutants did not show morphological differences with WT gemmae (Supplementary Fig. 4b). However, the gemmae of Mp*log* mutants showed upward growth of the thallus (Supplementary Fig. 4b, 5a). This defect of Mp*log* mutants was more substantial than that observed in KAI2-dependent signaling mutants. The application of BA rescued the defects (Supplementary Fig. 5b).

**Figure 3.**
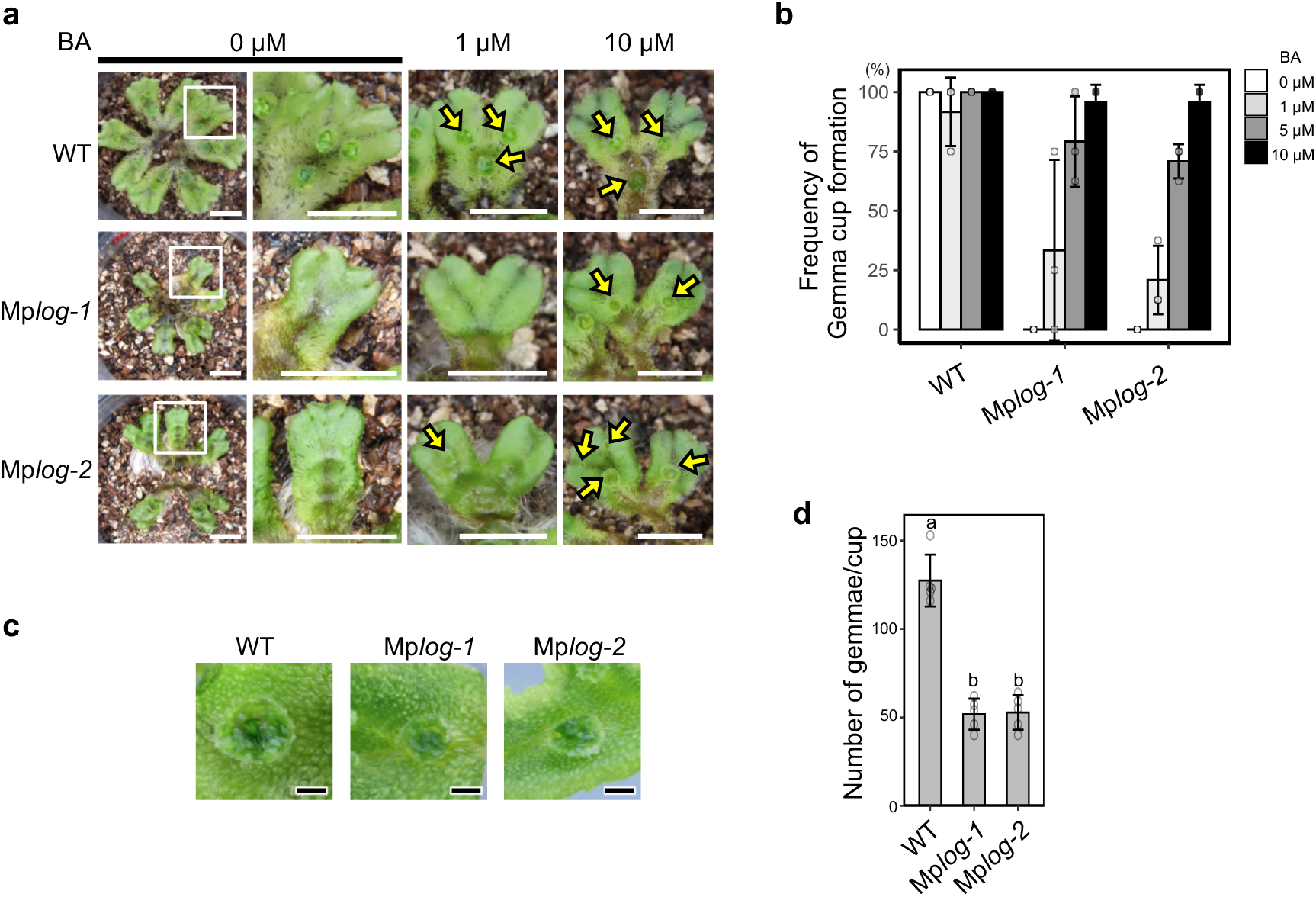
Cytokinins synthesized by Mp*LOG* control gemma cup and gemma formation. **a** Effects of cytokinin application on gemma cup formation in WT, Mp*log-1*, and Mp*log-2*. Plants were grown on vermiculite with 6-benzyl adenine (BA) application to the vermiculite once a week. Yellow arrows indicate gemma cups. Scale bars: 1cm. **b** Effect of cytokinin on the frequency of gemma cup formation in WT, Mp*log-1*, and Mp*log-2* plants. Plants were grown on 1/2 B5 medium for 6 days, then transferred to vermiculite and grown for 3 weeks or more with BA application to the vermiculite once a week. The frequency of gemma cup formation was scored at 8 points after the second bifurcation of the thallus tip. Bars represent mean ± SD (n=3). **c** The top view of gemma cups at day 14 stage in WT, Mp*log-1*, and Mp*log-2* plants grown on 1/2 B5 medium. Scale bars: 1 mm. **d** The number of gemmae in gemma cups at day 14 stage in WT, Mp*log-1*, and Mp*log-2* plants grown on 1/2 B5 medium. Bars represent mean ± SD (n=5). The Tukey’s HSD test was used for multiple comparisons and statistical differences (p<0.05) are indicated by different letters.

### Cytokinin works downstream of KAI2-dependent signaling

We showed that Mp*LOG* expression is positively regulated by KL signaling, and MpLOG catalyzes cytokinin synthesis. The defects in gemma cup and gemma formation observed in the Mp*log* mutants resembled those seen in KAI2-dependent signaling mutants, and Mp*log* defects were rescued by exogenous application of cytokinin (Fig. 3a, b). These results suggest that KAI2-dependent signaling controls gemma cup and gemma formation by controlling cytokinin synthesis. To test this hypothesis, we analyzed if defects in the Mp*kai2a* and Mp*max2* mutants are rescued by cytokinin. We found that defects in gemma cup formation in Mp*kai2a* and Mp*max2* were restored by addition of BA in a dose-dependent manner (Fig. 4a, b). The phenotype of upward growth of the thallus of Mp*kai2a* and Mp*max2* mutants was alleviated by the addition of BA in a dose-dependent manner (Supplementary Fig. 5c, d).

**Figure 4.**
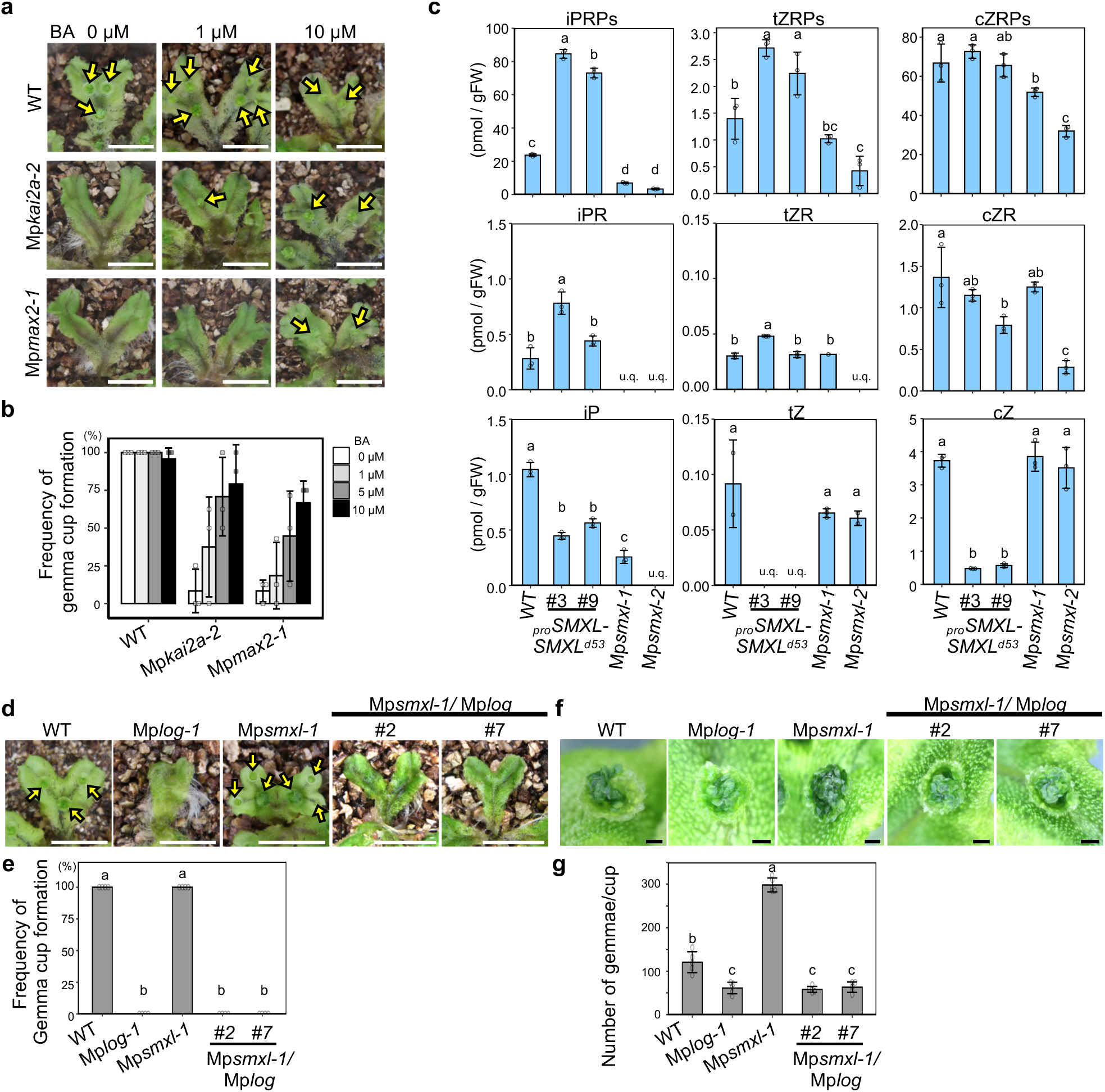
Cytokinin works downstream of KAI2-dependent signaling. **a** Effects of cytokinin application to gemma cup formation in WT, Mp*kai2a-2,* and Mp*max2-1.* Plants were grown on vermiculite with BA application to the vermiculite once a week. Yellow arrows indicate gemma cups. Scale bars: 1 cm. **b** Effect of cytokinin on the frequency of gemma cup formation in WT, Mp*kai2a-2,* and Mp*max2-1*. Plants were grown on 1/2 B5 medium for 6 days, then transferred to vermiculite and grown for 3 weeks or more with BA application to the vermiculite once a week. The frequency of gemma cup formation was scored at 8 points after the second bifurcation of the thallus tip. Bars represent mean ± SD (n=3). **c** Quantification of active cytokinins (iP, tZ, and cZ) in WT, Mp*smxl-1*, Mp*smxl-2, _pro_SMXL-SMXL^d53^*#3, and *_pro_SMXL-SMXL^d53^*#9 plants. Bars represent means ± SD (n=3). u.q. indicates under quantification limits. iPRPs, tZRPs, and cZRPs; ribotide precursors, iPR, tZR, and cZR; riboside precursors. **d, e** Genetic interaction between MpLOG and MpSMAXL in gemma cup formation. Thalli of WT, Mp*log-1*, Mp*smxl-1*, Mp*smxl-1*/Mp*log*#2, and Mp*smxl-1*/Mp*log*#7 plants grown on vermiculite (d). Yellow arrows indicate gemma cups. Scale bars: 1 cm. Quantification of gemma cup formation frequency (e). Bars represent mean ± SD (n=4). **f, g** Genetic interaction between MpLOG and MpSMAXL in gemma cup formation in the gemma number phenotype.Top view of day14 gemma cups of WT, Mp*log-1*, Mp*smxl-1*, Mp*smxl-1*/Mp*log*#2, and Mp*smxl-1*/Mp*log*#7 grown on 1/2 B5 medium (f). Scale bars: 1 mm. Quantification of the number of gemmae per cup (g). Bars represent the mean±SD (n=5). The Tukey’s HSD test was used for multiple comparisons and statistical differences (p<0.05) are indicated by different letters in (c), (e) and (f).

To further confirm that cytokinin acts downstream of KAI2-dependent signaling, which is dependent on the degradation of MpSMXL, we measured the contents of endogenous cytokinins (iP, tZ, cZ) in Mp*SMXL^d^*^53^overexpressing degradation-resistant MpSMXL, and in the Mp*smxl* loss of function mutants (Fig. 4c). The levels of cytokinins were decreased in the Mp*SMXL^d53^* mutants, in which KAI2-dependent signaling is disrupted. In particular, tZ declined to below quantification levels. On the other hand, the contents of tZ and cZ were not affected in Mp*smxl* mutants, for which the KAI2-dependent signaling is supposed to be enhanced. The level of iP was decreased in Mp*smxl* mutants for unknown reasons. Overall, our results support the hypothesis that KAI2-dependent signaling positively regulates cytokinin content.

Mp*smxl* loss-of-function mutants exhibit phenotypes that are opposite to those of Mp*kai2a* and Mp*max2*^11^. More gemmae per cup are formed in Mp*smxl*. To test the genetic relationship between Mp*SMXL* and Mp*LOG,* we generated Mp*smxl* Mp*log* double mutants. As expected if Mp*LOG* is downstream of MpSMXL, the Mp*smxl* mutant phenotypes were alleviated in the double mutants: the Mp*smxl* Mp*log* double mutants closely resembled the Mp*log* single mutants (Fig. 4d, f, Supplementary Fig. 5e). Gemma cups were normally formed in the Mp*smxl* mutants even grown on vermiculite, while it was severely suppressed in Mp*log* and in the double mutant (Fig. 4d, e). Mp*smxl* mutants produced an increased number of gemmae per cup when grown on medium, whereas this number was reduced in the Mp*smxl* Mp*log* double mutant (Fig. 4f, g), suggesting that the function of Mp*LOG* is required for the expression of the phenotypic traits exhibited by Mp*smx1*. Genetic analysis indicated that Mp*LOG* works downstream of the KAI2-dependent signaling pathway for gemma cup and gemma formation. In addition, the angle of the thallus relative to the horizontal line is smaller in Mp*smxl* and the thallus grew upward in the double mutant as in Mp*log* (Supplementary Fig. 5e, f), indicating that these traits are also under the control of Mp*LOG*.

### Control of MpLOG expression by KAI2-dependent signaling is cell-type specific

To understand the roles of cytokinins in the control of vegetative reproduction in *M. polymorpha*, we analyzed the spatial localization of Mp*LOG* expression. The 5 kb and 1 kb promoter regions of the Mp*LOG* gene were fused with the *EGFP* gene and a nuclear localization signal sequence (*_pro_*Mp*LOG*-EGFPNLS and *_pro_*Mp*LOG5kb*-EGFPNLS) and introduced into WT. Mp*kai2a* mutation was introduced into *_pro_*Mp*LOG*-EGFPNLS (*_pro_*Mp*LOG-EGFPNLS*/Mp*kai2a-7*). No difference was observed in the fluorescence pattern between the lines containing the 5 kb and 1 kb promoter regions (Fig. 5a, e, Supplementary Fig. 6). We used the plants containing *_pro_*Mp*LOG*-EGFPNLS with the 1 kb promoter region in the subsequent observations. EGFP fluorescence was observed in the meristem region at the notch and rhizoid initial cells of the gemma (Fig. 5a). The fluorescence in the meristem was not observed in the Mp*kai2a* mutant (Fig. 5b), indicating that the expression of Mp*LOG* in the meristem cells is under the control of KAI2-dependent signaling. In contrast, EGFP fluorescence in the rhizoid initial cells was observed in Mp*kai2a*. During gemma cup initiation, EGFP fluorescence was observed in the initiating cups in WT but not in Mp*kai2*, which is consistent with the absence of gemma cups in the Mp*kai2a* mutant (Fig, 5c, d). We further analyzed Mp*LOG* expression in the gemma cups and initiating gemmae using confocal microscopy. We observed that Mp*LOG* is expressed on the upper surface of the thallus in WT and Mp*kai2a*, indicating that the expression of Mp*LOG* in this region is not regulated by KAI2 signaling (Fig. 5e, f). Intense fluorescence was detected in the basal part of the gemma cup in WT (Fig. 5e). At the bottom of the gemma cup, fluorescence localized in gemma initial cells, initiating gemmae, slime papillae, and the gemma cup floor cells in WT (Fig. 5g). On the other hand, in Mp*kai2a* mutants grown on medium, EGFP fluorescence was observed in the slime papillae and gemma cup floor cells, but not in gemma initial cells, and initiating gemmae (Fig. 5h). Therefore, expression of Mp*LOG* in the meristem region in the gemmae, gemma initial cells and initiating gemmae is regulated by KAI2-dependent signaling while Mp*LOG* expression in the epidermis on the thallus, slime papillae, and gemma cup floor cells is not. We conclude that KAI2-dependent signaling controls gemma cup and gemma formation by enhancing cytokinin synthesis through the promotion of Mp*LOG* expression. Although their number is reduced, gemmae are formed in the Mp*kai2a* mutants when plants are grown on medium. However, Mp*LOG* expression was not observed in gemma initial cells and initiating gemmae in Mp*kai2a*. This suggests that either gemmae formed in the Mp*kai2a* mutants do not depend on Mp*LOG* or that residual expression levels of Mp*LOG* are sufficient for the generation of a small number of gemmae. It is also possible that the initiation of gemmae in Mp*kai2a* on the medium depends solely on nutrient conditions and is independent of cytokinin.

**Figure 5.**
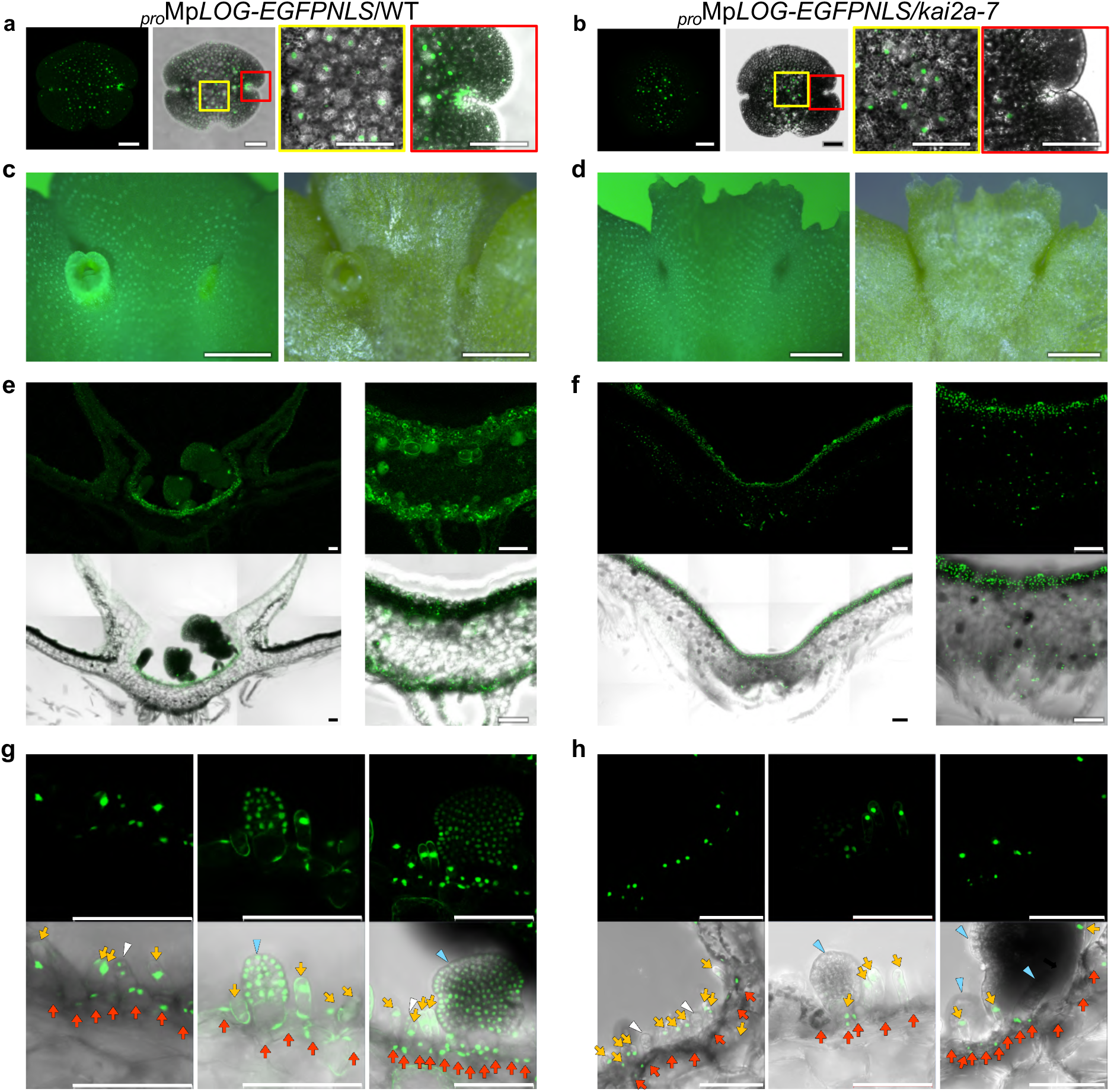
Control of Mp*LOG* expression by KAI2-dependent signaling. **a, b** Localization of EGFP fluorescence in the gemma of _pro_Mp*LOG-EGFPNLS/*WT (a), and _pro_Mp*LOG-EGFPNLS/kai2a-7* (b) grown on 1/2 B5 medium, observed by confocal laser scanning microscopy. From the left: EGFP fluorescence, EGFP merged with blight field image, enlargement of the region in yellow square. **c, d** Localization of EGFP fluorescence in the thallus of _pro_Mp*LOG-EGFPNLS/*WT (c), and _pro_Mp*LOG-EGFPNLS/kai2a-7* (d) grown on vermiculite, observed by stereo microscopy. Left: EGFP fluorescence, Right: blight field image. **e, f** Localization of EGFP fluorescence in the cross-sections of the midrib of _pro_Mp*LOG-EGFPNLS/*WT (e), and _pro_Mp*LOG-EGFPNLS/kai2a-7* (f) grown on vermiculite, observed by confocal laser scanning microscopy. Top panels: EGFP fluorescence, Bottom panels: EGFP merged with blight field image. **g, h** Localization of EGFP fluorescence in initiating gemmae, slime papilla, and gemma cup floor cells at the bottom of gemma cup in _pro_Mp*LOG-EGFPNLS/*WT (g), and _pro_Mp*LOG-EGFPNLS/kai2a-7* (h) grown on 1/2 B5 medium, observed by confocal laser scanning microscopy. White arrowheads: gemma initial cells, blue arrowheads: **i**nitiating gemmae, yellow arrows: slime papillae, red arrows: gemma cup floor cells. Scale bars: 100 µm.

### *GEMMA CUP-ASSOCIATED MYB1* (*GCAM1*) expression is promoted by cytokinin synthesized by MpLOG

*GCAM1* is an ortholog of *Arabidopsis REGULATOR OF AXILLARY MERISTEMS* (*RAX*) genes, encoding the R2R3-MYB transcription factors^47, 51^. *GCAM1* was first identified as the gene responsible for gemma cup formation. Subsequently, we demonstrated that *GCAM1* is also involved in gemma initiation and works downstream of KAI2-dependent signaling^11^. *GCAM1* expression is known to be positively regulated by cytokinin^52^. Hence, we postulated that *GCAM1* expression is downstream of cytokinin synthesis regulated by MpLOG. We first confirmed that *GCAM1* expression is enhanced by cytokinin application (Fig. 6a). In a prior study, we introduced *_pro_GCAM1-EGFPNLS,* containing approximately 5 kb of the of *GCAM1* promoter region, into WT plants. Fluorescence localized in gemma initial cells, initiating gemmae and slime papillae but not in gemma cup floor cells^11^. Here, we confirmed this expression pattern of *GCAM1* in WT plants (Fig. 6b). The localization of *GCAM1* expression observed in our studies differs slightly from the reported ones^47^. We believe that *GCAM1* expression in the gemma cup floor cells described in Yasui *et al.* (2019) is likely the signal in the slime papillae. EGFP fluorescence was absent from the gemma initial cells and initiating gemmae in the Mp*log* mutant (*_pro_*Mp*GCAM1-EGFPNLS*/Mp*log*), indicating that *GCAM1* expression in these cells depends on MpLOG activity (Fig. 6c). Comparisons of Mp*LOG* expression between WT and Mp*kai2a* mutants and *GCAM1* expression between WT and Mp*log* mutants support the idea that *GCAM1* expression in the gemma initial cells, initiating gemmae and possibly in slime papillae is promoted by cytokinins synthesized by MpLOG activity regulated by KAI2-dependent signaling (Fig. 7a).

**Figure 6.**
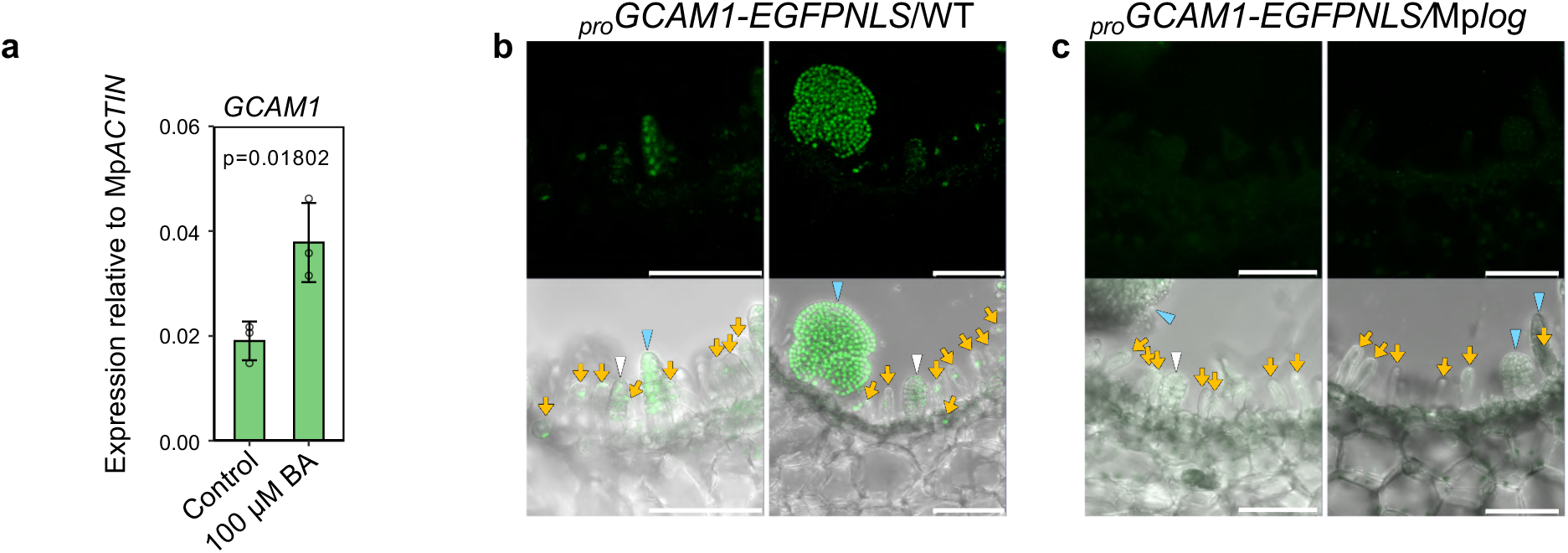
*GEMMA CUP-ASSOCIATED MYB1 (GCAM1)* expression is promoted by cytokinin synthesized by MpLOG. **a** The induction of *GCAM* expression by cytokinin was examined using qPCR analysis. Expression levels relative to Mp*ACTIN* are shown. Bars represent means ± SD (n=3). The p values were calculated by using the Student’s t-tests. **b, c** Localization of EGFP fluorescence in initiating gemmae at the bottom of gemma cup in *_pro_GCAM1-EGFPNLS/*WT (b) and *_pro_GCAM1-EGFPNLS/*Mp*log* (c) grown on 1/2 B5 medium and observed by confocal laser scanning microscopy. White arrowheads: gemma initial cells, blue arrowheads: **i**nitiating gemmae, yellow arrows: slime papillae. Scale bars: 100 µm.

**Figure 7.**
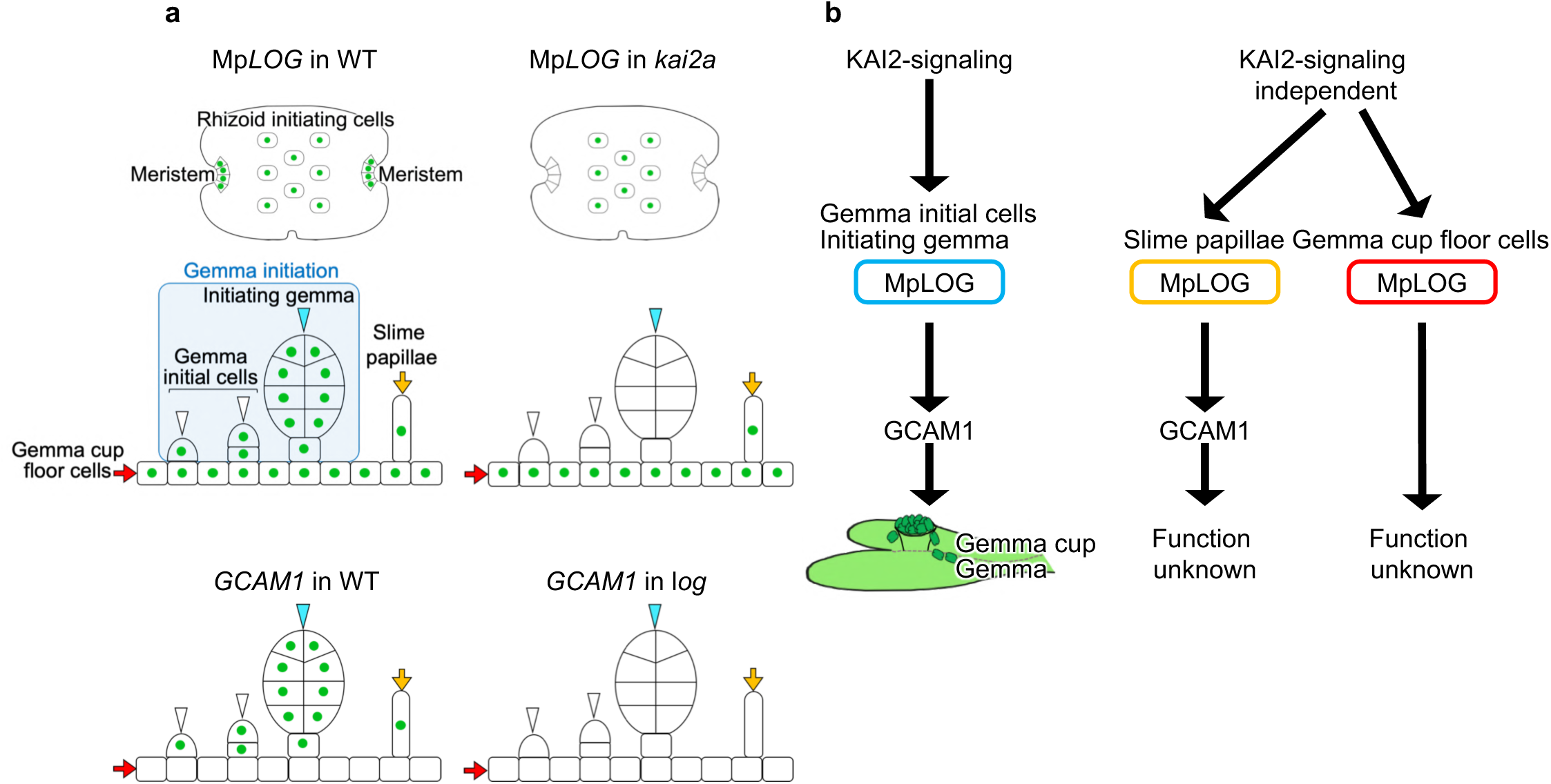
Model of cytokinin-mediated control of gemma cup and gemma formation by KAI2-dependent signaling in *M. polymorpha*. **a** Schematic diagram of the Mp*LOG* and *GCAM1* expression in WT, Mp*kai2a*, and Mp*log* mutant plants. Green circles indicate EGFP fluorescence. White arrowheads: gemma initial cells, blue arrowheads: initiating gemmae, yellow arrows: slime papillae, red arrows: gemma cup floor cells. **b** The model for gemma cup and gemma formation by KAI2-dependent signaling. The activation of Mp*LOG* expression by KAI2-dependent signaling in gemma initial cells and initiating gemmae leads to the induction of *GCAM1* expression, resulting in gemma-cup and gemma formation. Mp*LOG* expression in slime papillae and gemma cup floor cells is KAI2 signaling independent. Mp*LOG* expression induces *GCAM1* expression in slime papillae but not in gemma cup floor cells. The functional significance of GCAM1 in slime papillae and MpLOG in gemma cup floor cells is unknown.

## Discussion

In this study, we demonstrated that KAI2-dependent signaling positively controls the expression of Mp*LOG*, leading to increased cytokinin levels. Defects in the Mp*log* mutants resembled those of KAI2-dependent signaling mutants. In addition, Mp*log* and KAI2-dependent signaling mutants were rescued by the exogenous supply of cytokinin, supporting the idea that cytokinin synthesis by MpLOG works as a downstream regulator of KAI2-dependent signaling. We also showed that *GCAM1*, the primary regulator of gemma cup and gemma formation, is activated by cytokinin produced by MpLOG. We propose a genetic cascade, starting from KAI2-dependent signaling, leading to vegetative reproduction through the induction of Mp*LOG* and *GCAM1* (Fig. 7b).

Plant growth and development rely on the continuous activity of the shoot apical meristem (SAM), where stem cell proliferation promoted by cytokinin is pivotal^53^. *LOG*s genes, expressed in the stem cell zone of the SAM, play crucial roles in maintaining the stem cell population in the SAM^46^. In *Arabidopsis*, cytokinins synthesized in the tip of the meristem, facilitated by specific localization of *LOG4* expression, serves as the source of a gradient of cytokinin activity, which acts as the spatial cue in the SAM^54^. *LOG* expressed at the tip of meristems controls stem cell proliferation in rice panicle^46^. We showed that Mp*LOG* is expressed in the meristem region, gemma initial cells, and initiating gemmae that contain undifferentiated meristematic cells. The LOG activity localized in these regions likely contributes to the initiation and growth of gemma cups and gemmae, facilitating vegetative reproduction. Spatially regulated *LOG* expression may be crucial for the proliferation of undifferentiated cells during vegetative reproduction in *M. polymorpha*. This indicates a possibility that the function of *LOG* as a regulator of the undifferentiated cells was established in the common ancestor of land plants. Considering that adenosine phosphate-IPTs and CYP735A are of later origin, the specification of undifferentiated meristematic cells by localized cytokinin synthesis achieved by spatiotemporal regulation of LOG activity might be an ancient and fundamental mechanism^55, 56, 57^. LOG genes are present in all major lineages of prokaryotes and plants^58^. LOG-containing microbial pathogens cause infection in a broad range of hosts, including plants, insects, and animals, implying the importance of cytokinins as communication molecules in cross-kingdom biological interactions. In *Mycobacterium tuberculosis*, which exclusively infects animal cells, the LOG protein levels are tightly controlled through 26S proteasome-dependent degradation, possibly to fine-tune cytokinin production^59^. The control of cytokinin synthesis by LOG may be a more universal and widespread system than previously thought and achieved in diverse ways. Further studies on LOG’s functions, diversity, and evolution will uncover new insights into the role of cytokinins.

Both gemma cup formation and gemma formation are critical steps for vegetative reproduction; they are distinct developmental processes that occur in different cells at different developmental stages^7^. We previously reported that GCAM1 promotes gemma formation in addition to gemma cup formation^11, 47^. In this study, we showed that the KAI2 signaling-LOG module controls these two steps by activating GCAM1. The control of cytokinin synthesis by KAI2-dependent signaling is highly cell-type specific (Fig. 7a, b). In the gemma, Mp*LOG* is expressed in the meristem region and the rhizoid initial cells; however, the expression in the meristem region but not the rhizoid initial cells is KAI2 signaling-dependent. Similarly, among Mp*LOG* expression in four types of cells at the bottom of the gemma cup: gemma initial cells, initiating gemmae, slime papillae, and gemma cup floor cells, expression in slime papillae and floor cells is KAI2 signaling-independent. This indicates that KAI2-dependent signaling is responsible for only a subset of Mp*LOG* expression (Fig. 7a). Coincidentally, *GCAM1* is expressed in cells where Mp*LOG* is activated by KAI2-dependent signaling, suggesting that KL signaling may be profoundly involved in controlling vegetative reproduction. The underlying mechanisms for the cell-type specificity of KAI2-dependent signaling are unknown. Considering that the receptor MpKAI2A localizes in wider regions, a possible explanation is that the cell type-specificity may be regulated at the level of KL biosynthesis^11^.

*GCAM1* orthologs control axillary meristem formation in *Arabidopsis*, tomato, and pepper^51, 60, 61,62^. This suggests a deep homology between shoot branching in angiosperms and gemma formation in *M. polymorpha*. However, although cytokinin is the upstream regulator of *GCAM1*, *RAX* genes work upstream of cytokinin and promote axillary meristem formation by enhancing cytokinin synthesis and signaling. We further demonstrated that the cytokinin-GCAM1 module is controlled by KAI2-dependent signaling in *M. polymorpha*. However, control of shoot branching by KAI2-dependent signaling is not known, and the cross-regulation of KAI2 signaling and cytokinin has not been demonstrated in angiosperms. In angiosperms, strigolactones control shoot branching at the step of axillary bud outgrowth, through interaction with cytokinin and auxin^63, 64, 65^. Strigolactones acquired the role of a class of plant hormones by gene duplication of KAI2 in seed plants^39, 41, 66, 67^. Subsequently, the increase in the number and functional diversification of SMXL genes brought about the specificity of the two signaling pathways^67^. An intriguing possibility is that the link between KAI2-dependent signaling and cytokinin was taken over by strigolactone signaling in angiosperms. We cannot rule out the possibility that KAI2 signaling-cytokinin cross-regulation was acquired in the liverwort lineage.

Cytokinins are present in all land plants, algae, and some fungi and bacteria. Cytokinin biosynthesis and signaling pathways in plants have been mainly studied in angiosperms, however, some reports can be found outside angiosperms^48, 68, 69^. In angiosperms, cytokinins are synthesized through adenosine phosphate-IPTs and tRNA-IPT. Phylogenetic analysis revealed that adenosine phosphate-IPTs are derived from tRNA-IPT^55, 56^. Although the exact timing remains controversial, the duplication of tRNA-IPT leading to the adenosine phosphate-IPTs occurred later than the divergence of bryophytes^55, 56^. Currently, the mechanism of cytokinin biosynthesis in bryophytes is not known, as cognate orthologs of angiosperm adenosine phosphate-IPTs are not present^57^. Bryophytes do not contain orthologs of CYP735A either^37, 70^. The function of LOG is also enigmatic. Severe reduction of active cytokinins and extensive increases in their precursors in Mp*log* mutants support the conserved function of LOG in *M. polymorpha*. However, despite a single *LOG* gene in the genome of *M. polymorpha,* active cytokinins are still present in complete loss of function mutants of Mp*LOG*. This suggests the presence of LOG-independent pathways of cytokinin biosynthesis in *M. polymorpha*. Angiosperms contain a small family of LOG genes, but cytokinin levels in plants that lost the function of all *LOG* genes have yet to be reported^71^. Therefore, the generality of the LOG-independent cytokinin synthesis pathway is unclear. Recently, the cytokinin/purine riboside nucleosidase1 (CPN1), which catalyzes the deribosylation of cytokinin nucleoside precursors and other purine nucleosides, was discovered in rice, supporting the existence of LOG-independent cytokinin synthesis^72^. The ortholog of *CPN1* exists in hornworts, the basal lineage of bryophytes, but not in *M. polymorpha* and *P. patens*, indicating that *CPN1* occurred in the common ancestor of land plants but was lost in *M. polymorpha* and *P. patens*. While CPN1 contains double Ribo hydro-like domains, *CPN1* homologs with the single Ribo hydro-like domain of unknown function exist in land plants, including *M. polymorpha* (Supplementary Fig. 7). Further understanding of cytokinin synthesis in land plants in general and *M. polymorpha* specifically will help better understand the significance of KL-cytokinin cross-regulation in controlling vegetative reproduction and plant evolution.

## Methods

### Plant Materials and Growth Conditions

Strain Takaragaike-1 (Tak-1; Japanese male line) was used as the *M. polymorpha* wild-type (WT) in this study. Mp*AtD14ox*, Mp*kai2a-2*, Mp*max2-1*, Mp*smxl-1*, Mp*smxl-2*, *proSMXL-SMXL^d53^#9*, *proSMXL-SMXL^d53^#3*, and *proGCAM1-EGFPNLS* were described previously^11, 12^. Plants were generally cultured on half-strength Gamborg’s B5 medium with 1% agar, under continuous light (50 ∼ 60 µmol photons m⁻²s⁻¹) at 22°C.

### RNA extraction and expression analysis

RNA extraction, cDNA synthesis, and quantitative PCR were performed as previously described^11,12^. Primers used are listed in Table S3.

### RNA-seq analysis

Gemmae of Mp*AtD14ox* were incubated on growth medium for 24 hours at 22 °C under continuous light before being transferred to a liquid medium containing either acetone as the control or 1 µM *rac*-GR24, and then incubated for 3 or 6 hours. Total RNA was extracted from the plants using the NucleoSpin RNA Plant Kit (Macherey-Nagel). Seven biological replicates were prepared for each condition. Total RNA extraction, mRNA isolation, and RNA-seq library synthesis were performed using the MGISEQ-2000 sequencing platform (MGI Tech). The resultant reads were mapped to the *M. polymorpha* genome version 5.1r1 assembly using HISAT2(v2.1.0) with the default parameters. Mapped reads were counted using the featureCounts function of the Rsubread package (v2.2.6) in R. Gene expression between the samples treated with acetone- and 1 µM *rac*-GR24 was compared using DESeq2^73^. Genes with a *padj* < 0.05 were considered differentially expressed. GO enrichment analysis was conducted as previously described^3, 74^. The groups of genes that were upregulated or downregulated in the 3-hour and 6-hour treatments are summarized in Supplementary Table 1.

### Mutagenesis by CRISPR

CRISPR/Cas9-based genome editing of Mp*LOG* was performed as previously described^75^. The vector, pMpGE011-MpLOG containing the target sequences (gRNA1 and gRNA2 of Mp*LOG*), was introduced into WT. The vector including gRNA1 of Mp*LOG* was also introduced into Mp*smxl-1* strain. Respective target sites, genomic sequence changes, and amino acid sequence changes of Mp*log-1* and Mp*log-2* are shown in Supplementary Figure 4a. About Mp*smxl-1/*Mp*log* #2 and #7, genomic sequence changes, and amino acid sequence changes are shown in Supplementary Table 2. pMpGE010-MpKAI2A-gRNA1 described previously^75^ was introduced into the *_pro_*Mp*LOG-EGFPNLS* strain described below and *_pro_*Mp*LOG-EGFPNLS*/Mp*kai2a-7* was selected.

### Quantification of cytokinins

WT, Mp*log-1*, Mp*log-2*, Mp*smxl-1*, Mp*smxl-2*, *proSMXL-SMXL^d53^#9*, and *proSMXL-SMXL^d53^#3* were sown on growth medium and cultivated for seven to ten days. Approximately 100 mg of fresh weight of each plant was used for the hormone extraction. Analysis of cytokinins was performed as previously described^50, 76^. Approximately 100 mg of each sample was used for measurement of cytokinins by ultra-performance liquid chromatography (UPLC)-tandem mass spectrometry (AQUITY UPLC System/XEVO-TQXS; Waters, USA) with an ODScolumn (AQUITY UPLC HSS T3, 1.8 mm, 2.1 x 100 mm; Waters).

### Phenotypic analysis

Analysis of the thallus phenotypes, gemma-cup formation, and the number of gemmae in a gemma-cup were performed as described previously^11^. For the comparison of the gemma formation among WT, Mp*log-1*, Mp*smxl-1*, Mp*smxl-1*/Mp*log#2*, and Mp*smxl-1*/Mp*log#7,* the apical parts, which had not yet bifurcated and formed a gemma cup, were incubated on new growth medium for 14 days. Gemmae of WT, Mp*log-1*, Mp*smxl-1*, Mp*smxl-1*/Mp*log#2*, and Mp*smxl-1*/Mp*log#7* were transferred to vermiculite after 6 days of incubation on growth medium and observed from the side after another 14 days of incubation.

### Promoter-reporter analysis

About 5 kb and 1 kb of the promoter regions of Mp*LOG* were amplified from WT genomic DNA by PCR using the primer pairs, proMpLOG-5kb_F/proMpLOG_R, and proMpLOG1kb_F/proMpLOG_R, and cloned into pENTR/D-TOPO (Thermo Fisher Scientific). The generated entry vectors, pENTR-*_pro_*Mp*LOG5kb*, and pENTR-*_pro_*Mp*LOG* were transferred into the binary vectors pMpGWB315-EGFPmodified. The transformants generated by using the resulting vectors, pMpGWB315-EGFPmodified-*_pro_*Mp*LOG5kb* and pMpGWB315-EGFPmodified-*_pro_*Mp*LOG,* were observed as described previously^11^.

### Quantification and statistical analysis

Statistical analyses were performed in R version 4.1.2 (R Development Core Team) using the multcomp R package. Significant differences (p<0.05) were tested using Tukey’s honestly significant difference (HSD) test for multiple comparisons. Data are presented as the mean ± SD.

### Data availability

Raw RNA sequencing datasets generated in this study were deposited to Short Read Archive at DNA Data Bank of Japan (DDBJ) under DRA019200 (BioProject: PRJDB18746). Source Data for Fig. 1d-g, 2b, 3b, 3d, 4b-c, 4e, 4g, 6a, and Supplementary Fig. 2a-b, 5b, 5d, 5f are provided with this paper.

## Acknowledgments

We thank Dr. Yoshiya Seto and Dr. Xiaonan Xie for providing (–)-GR24. This research was supported by a Grants-in-Aid from the Ministry of Education, Culture, Sports, Science, and Technology, Japan (23H05409, 20H05684, and 17H06475 to J.K., JP21K15116 to A.K.) and Canon Foundation to J.K.

## Author contributions

J.K. designed the research and wrote the article. A.K., M.F., K. K., H.S., and Y.H. produced mutants, analyzed phenotypes, and examined gene expression. Y.T., M.K., and H.S. quantified cytokinins.

---

**Supplementary Figure 1.**
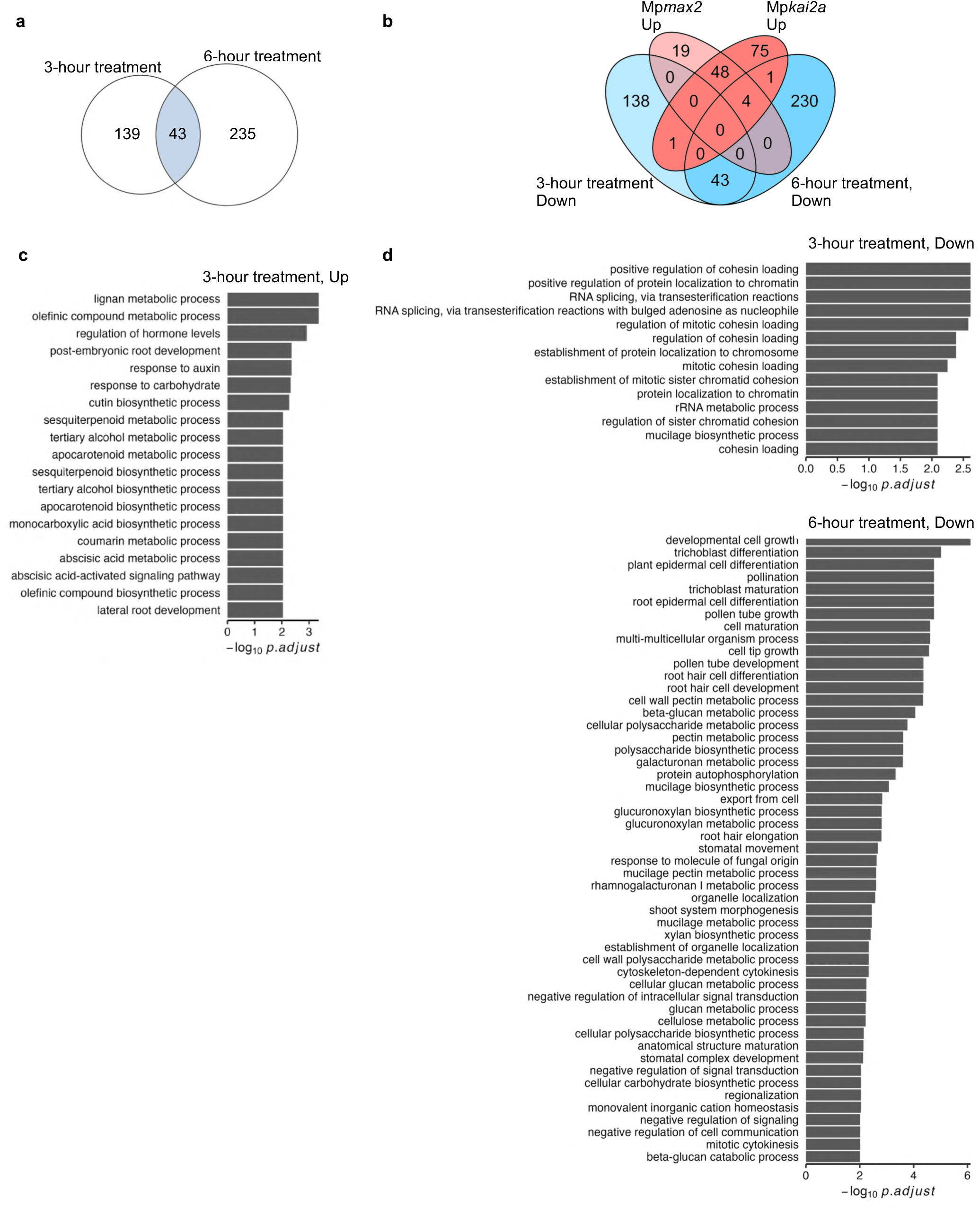
RNA-seq analysis of *rac*-GR24-treated plants compared to control plants after 3 or 6 hours. **a** Venn diagram showing the number of down-regulated genes after 1 µM *rac-*GR24 treatment for 3 and 6 hours. **b** Venn diagram showing the number of genes down-regulated genes after 1 µM *rac-* GR24 treatment, and up-regulated in Mp*max2-2* and Mp*kai2a-1* mutants. **c** Gene ontology (GO) analysis of the up-regulated genes after 3 hours of treatment. **d** GO analyses of the down-regulated genes after 3 hours (top) and 6 hour of treatment (bottom).

**Supplementary Figure 2.**
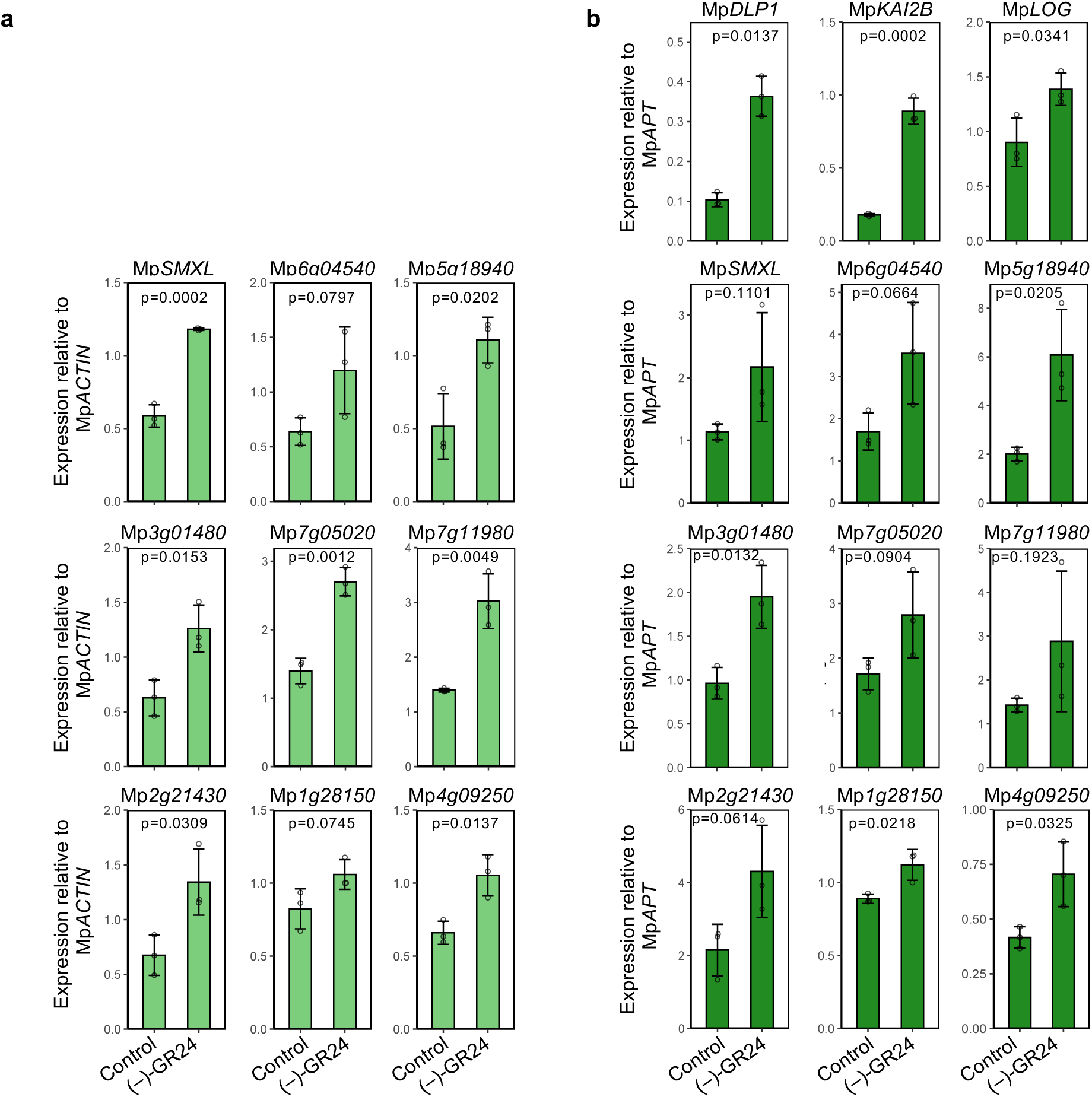
Expression levels of genes controlled by KAI2-dependent signaling. **a** Induction of 9 up-regulated genes by treatment with (–)-GR24 at 1 µM for 6 hours examined by quantitative PCR (qPCR) analysis in WT. Mp*ACTIN* was used as a reference gene. **b** Induction of 12 up-regulated genes by treatment with (–)-GR24 at 1 µM for 6 hours examined by quantitative PCR (qPCR) analysis in WT. *ADENINE PHOSPHORIBOSYL TRANSFERASE* gene (Mp*APT)* was used as a reference gene. The p values were calculated by using the Student’s t-tests.

**Supplementary Figure 3.**
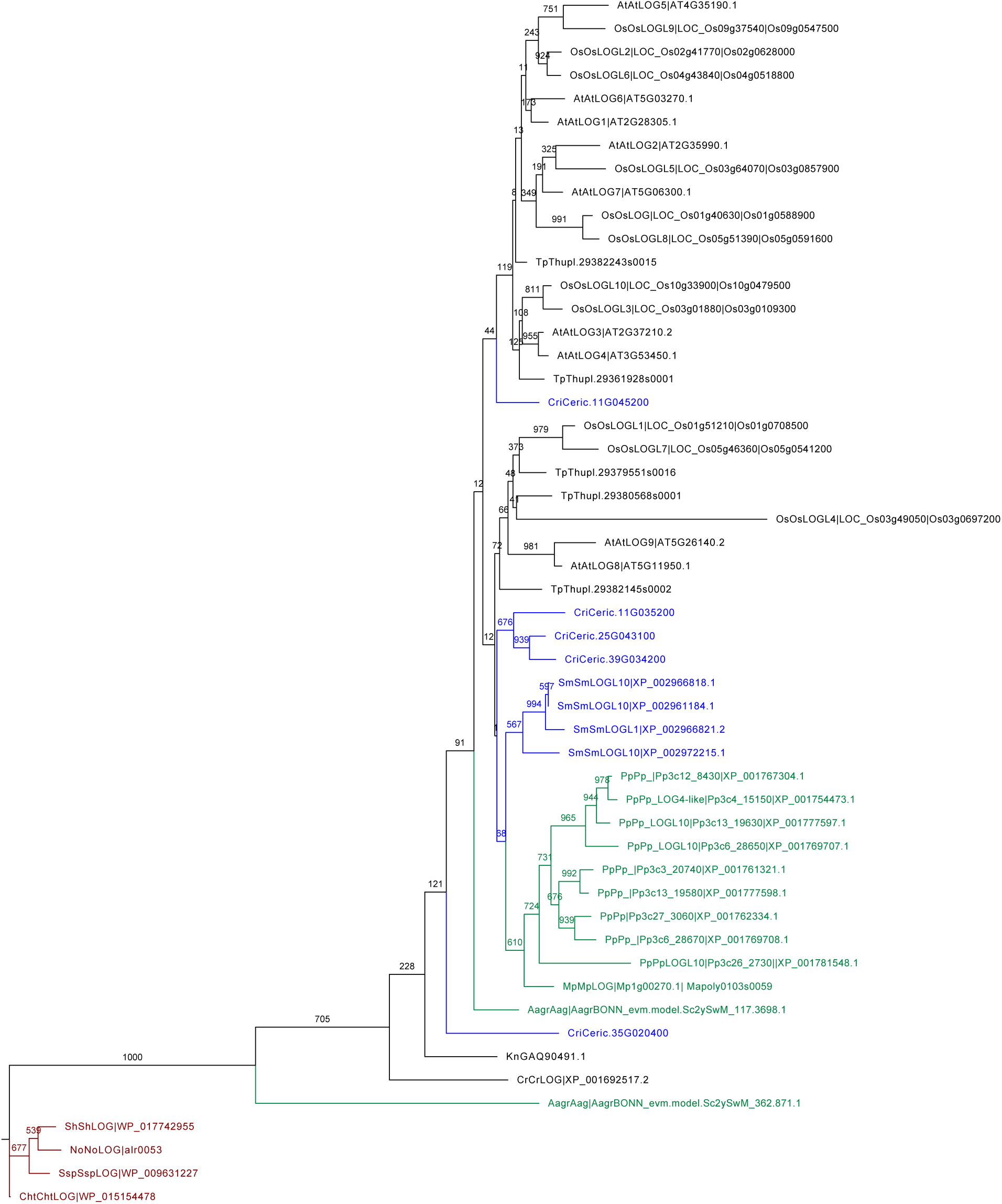
Phylogenetic tree of *LOG* genes. The phylogenetic analysis was performed using PhyML (http://www.atgc-montpellier.fr/phyml/), based on the maximum likelihood method with 1000 times bootstrap. Branches are colored according to plant lineages. Terminals are labeled with the abbreviations of the species name and gene IDs. Os, *Oryza sativa*; At, *Arabidopsis thaliana*; Cri, *Ceratopteris richardii*; Sm, *Selaginella moellendorffii*; Pp, *Physcomitrium patens*; Mp, *Marchantia polymorpha*; Aag, *Anthoceros agrestis*; Kn, *Klebsormidium nitens*; Cr, *Chlamydomonas reinhardtiis*; Sh, *Scytonema hofmannii*; No, Nostoc; Ssp, Synechocystis; Cht, Chroococcidiopsis. The bootstrap values are indicated at the branch point.

**Supplementary Figure 4.**
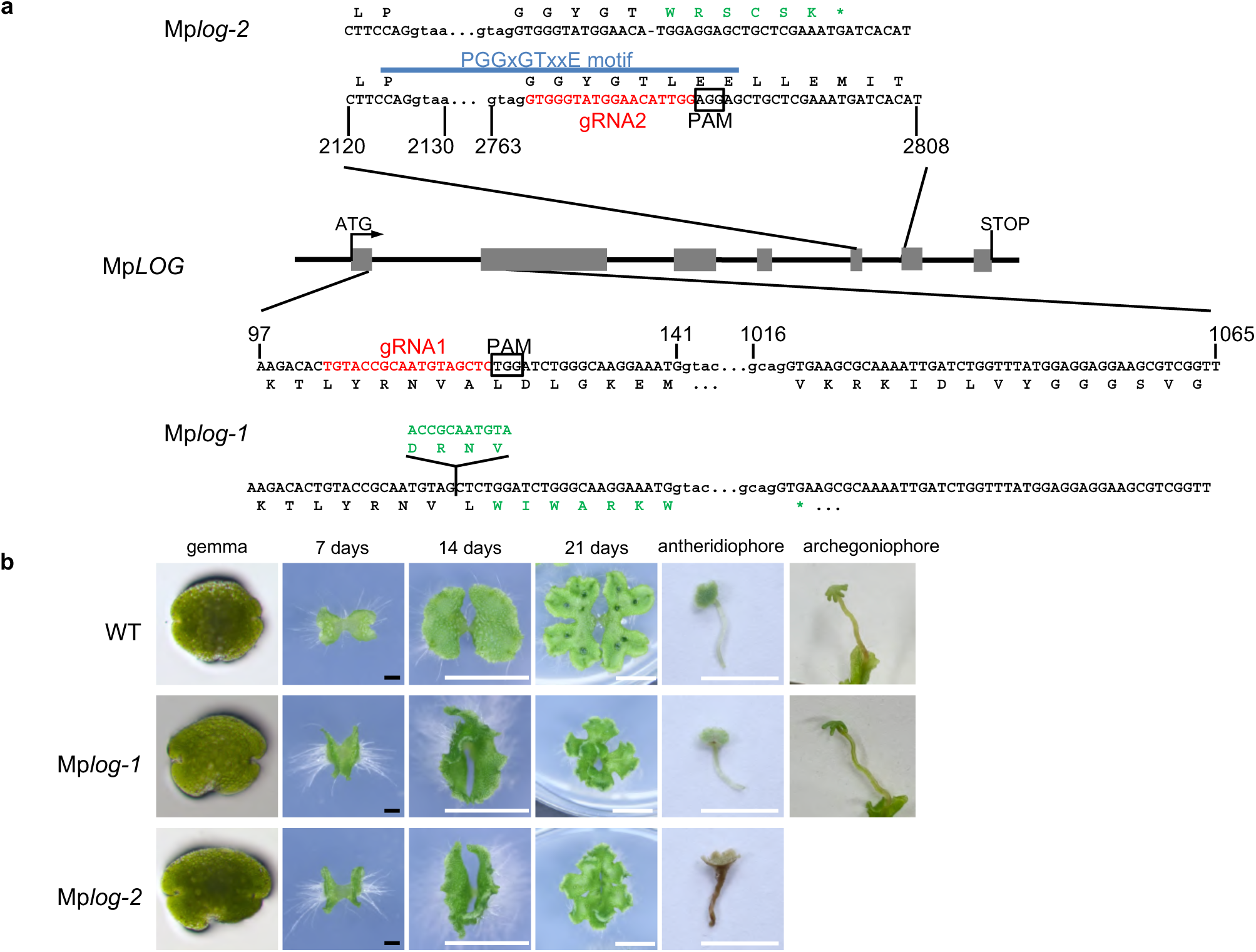
Production of Mp*log* loss of function mutants by CRISPR and their phenotypes. **a** Structure of the Mp*LOG* gene, the position of guide RNAs, PAM sequences, and resultant mutations are shown. The predicted amino acid sequences of the wild type and mutant LOG protein are shown in black and green letters, respectively. Asterisks indicate stop codons. **b** Morphological phenotypes of Mplog mutants, Mp*log-1,* Mp*log-2.*Thalli of Mp*log* mutants are smaller than those of WT and grow upwards. Sexual organs do not show obvious morphological defects and produce spores normally. Scale bars: 1 cm (white), 1 mm (black).

**Supplementary Figure 5.**
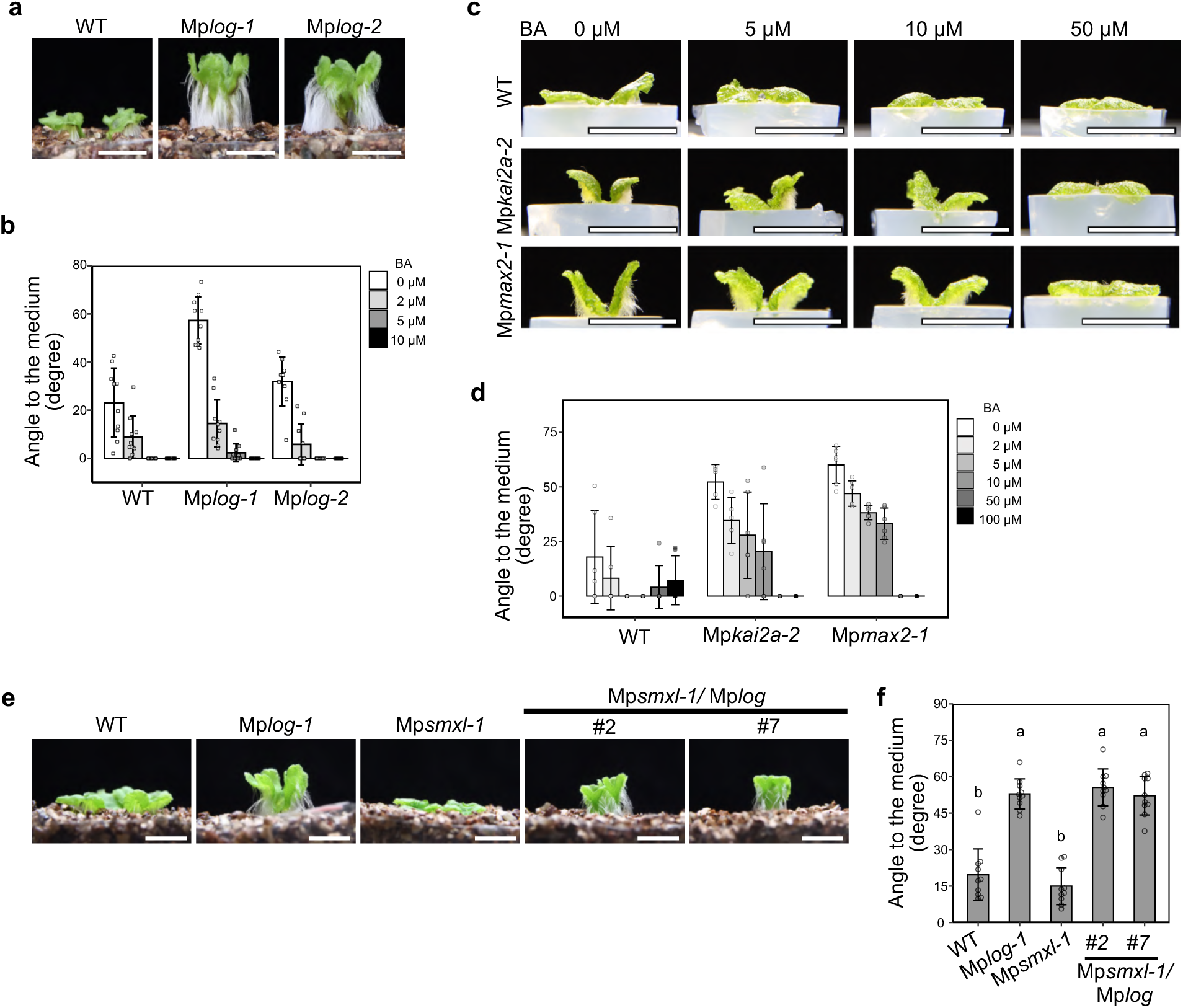
Cytokinin positively controls the flat growth of thallus. **a** Side view of 14-day-old thalli in WT, Mp*log-1*, and Mp*log-2* grown on vermiculite. **b** Effect of cytokinin on thallus angle to horizontal in WT, Mp*log-1*, and Mp*log-2.* Plants were grown on 1/2 B5 medium containing 0, 2, 5, or 10 µM 6-benzyl adenine (BA). Data are the mean ± SD (n=10). **c** Side view of 14-day-old thalli in WT, *kai2a-2*, and Mp*max2-1*. Plants were grown on 1/2 B5 medium containing 0, 5, 10 or 50 µM BA. **d** Effect of cytokinin on thallus angle to horizontal in WT, Mp*kai2a-2*, and Mp*max2-1* grown on 1/2 B5 medium containing 0, 2, 5, 10, 50 or 100 µM BA. Data are the mean ± SD (n=6). **e** Side view of thalli in WT, Mp*log-1*, Mp*smxl-1*, Mp*smxl-1*/Mp*log*#2, and Mp*smxl-1*/Mp*log*#7 grown on vermiculite. **f** Angles between the 14-d-old thalli and the growth media in WT, Mp*log-1*, Mp*smxl-1*, Mp*smxl-1*/Mp*log*#2, and Mp*smxl-1*/Mp*log*#7grown on 1/2 B5 medium. The Tukey’s HSD test was used for multiple comparisons and statistical differences (p<0.05) are indicated by different letters. Scale bars: 1 cm.

**Supplementary Figure 6.**
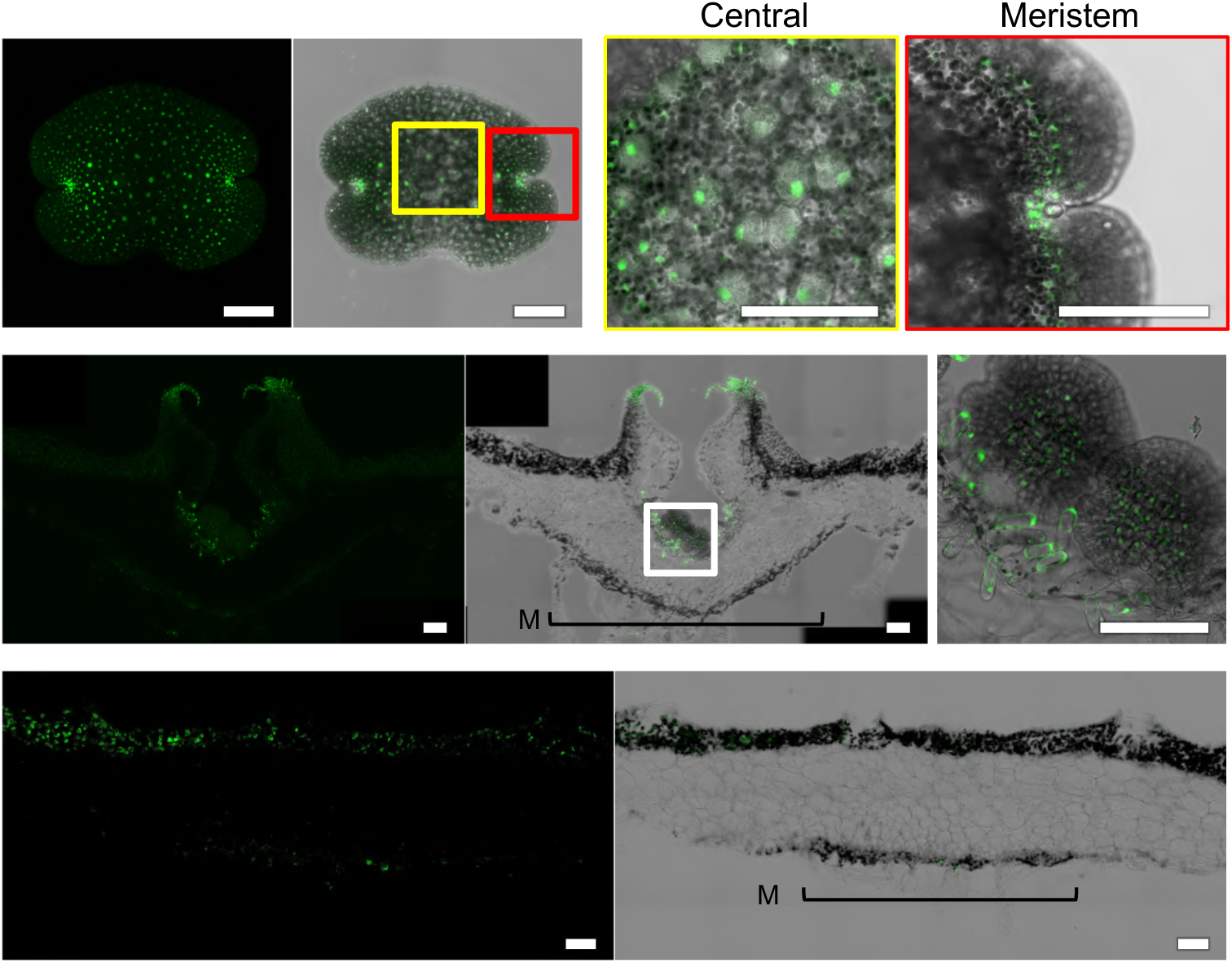
The expression pattern of Mp*LOG*. Localization of EGFP fluorescence in gemma (top), in the cross-sections of the gemma cup (middle) and in the midrib (bottom) of *_pro_*Mp*LOG5kb-EGFPNLS* containing the 5 kb region from the translation start codon. M: midrib region. Scale bars: 100 µm.

**Supplementary Figure 7.**
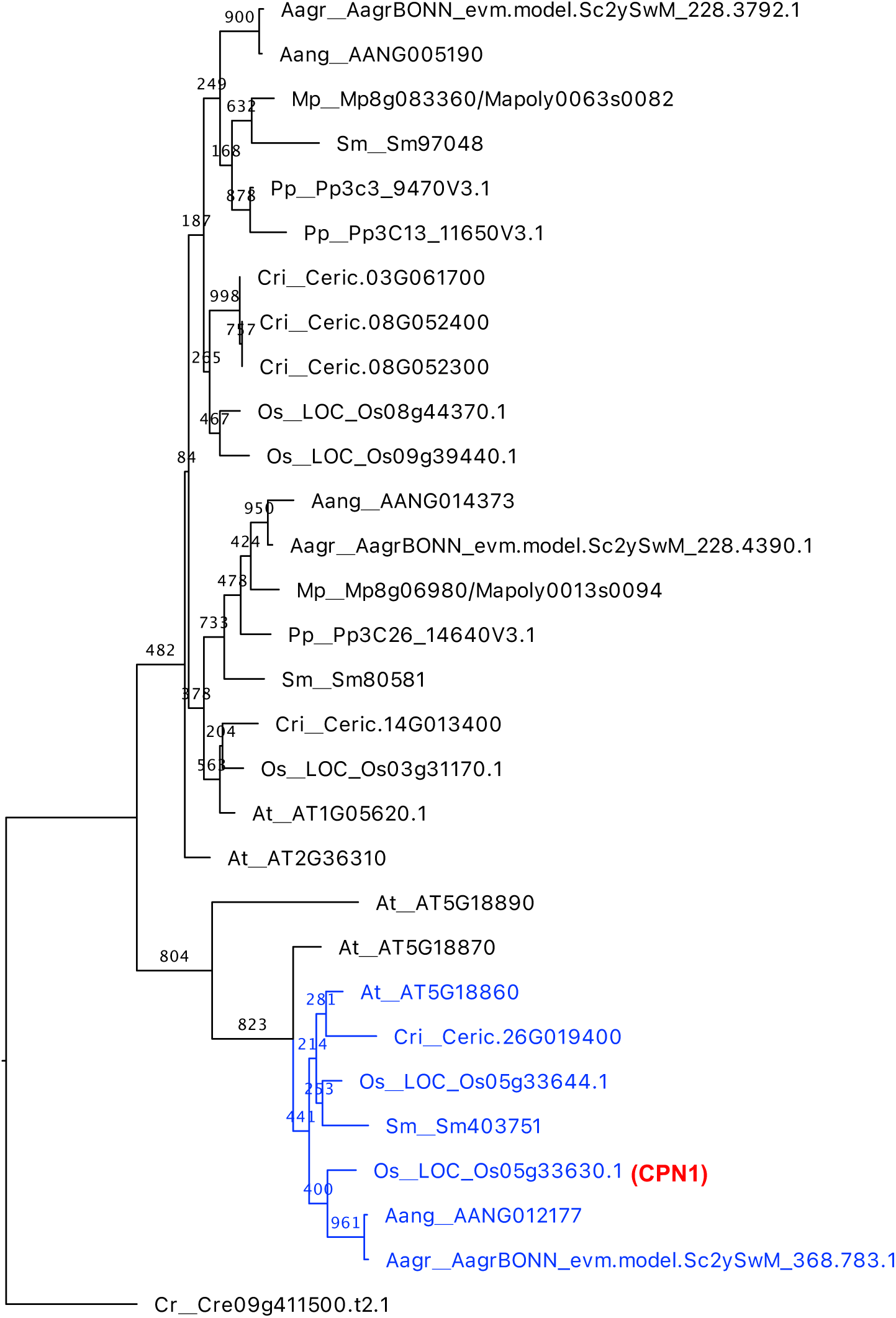
Phylogenetic tree of *CPN1* genes. The phylogenetic analysis was performed using PhyML (http://www.atgc-montpellier.fr/phyml/), based on the maximum likelihood method with 1000 times bootstrap. Terminals are labeled with the abbreviations of the species name and transcript ID. Os, *Oryza sativa*; At, *Arabidopsis thaliana*; Cri, *Ceratopteris richardii*; Sm, *Selaginella moellendorffii*; Pp, *Physcomitrium patens*; Mp, *Marchantia polymorpha*; Aagr, *Anthoceros agrestis*; Aang, *Anthoceros angustus*. Genes containing double tandem Ribo hydro-like domains are shown in blue. The bootstrap values are indicated at the branch point.

**Supplementary Table 1.**
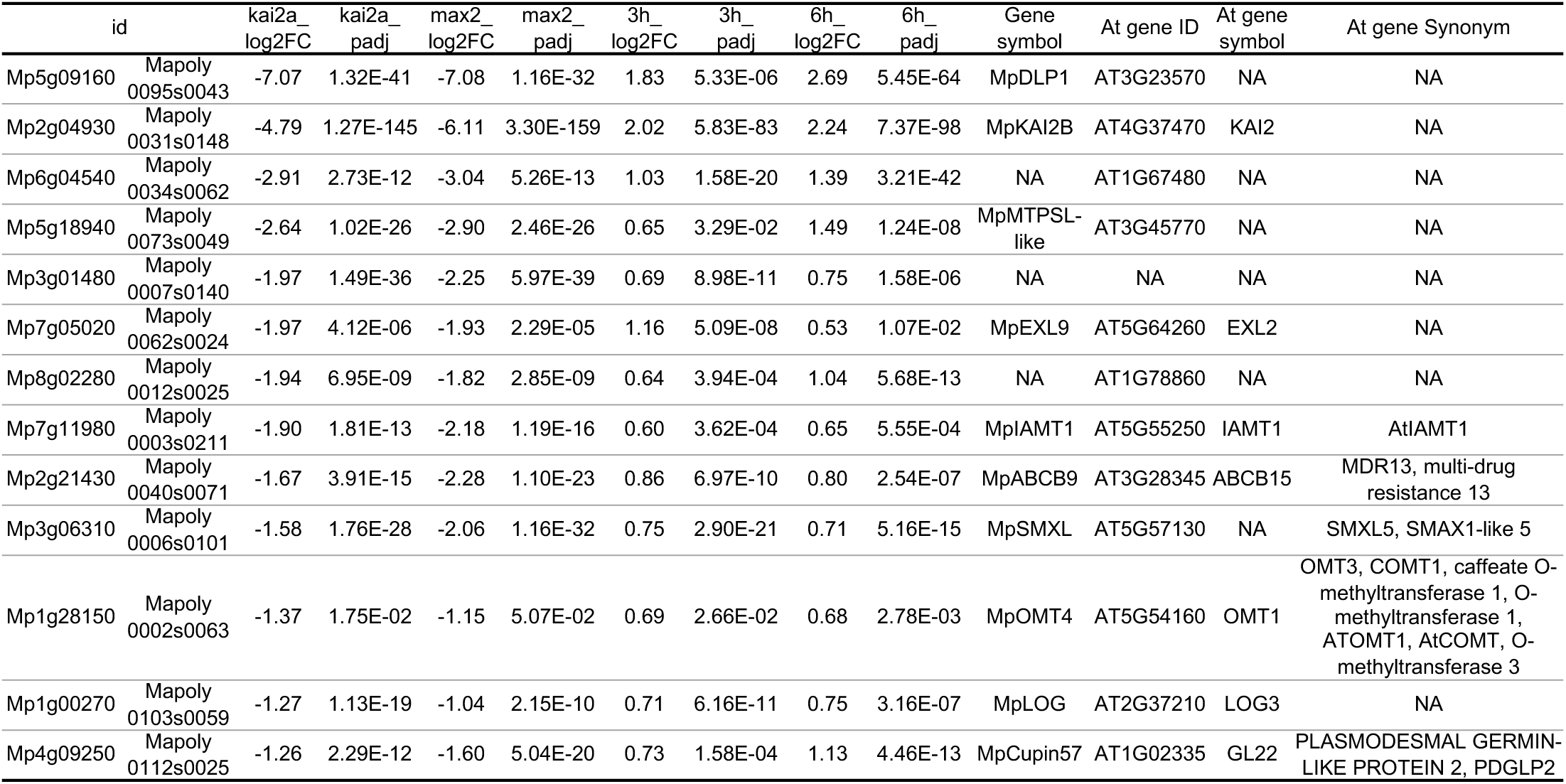
List of genes up-regulated in 3-hour and 6-hour treatment, and down-regulated in Mp*kai2a* and Mp*max2*.

**Supplementary Table 2.**
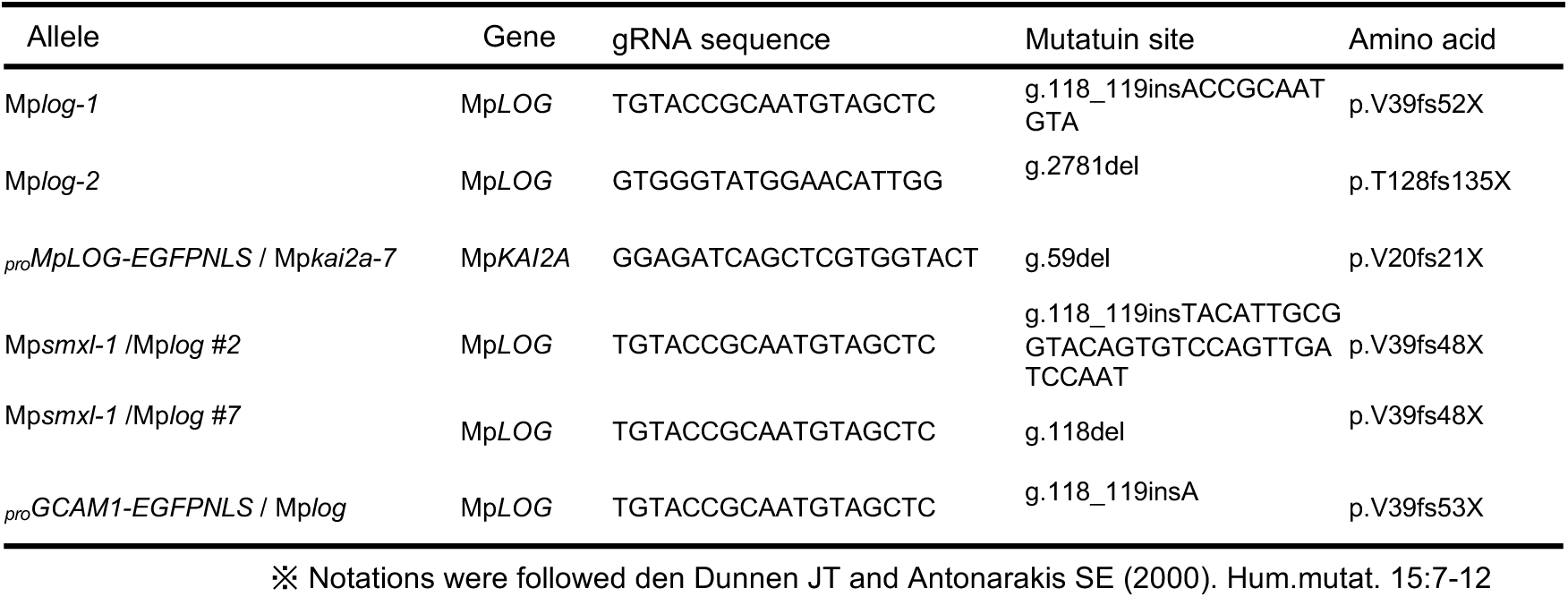
Production of CRISPR mutants.

**Supplementary Table 3.**
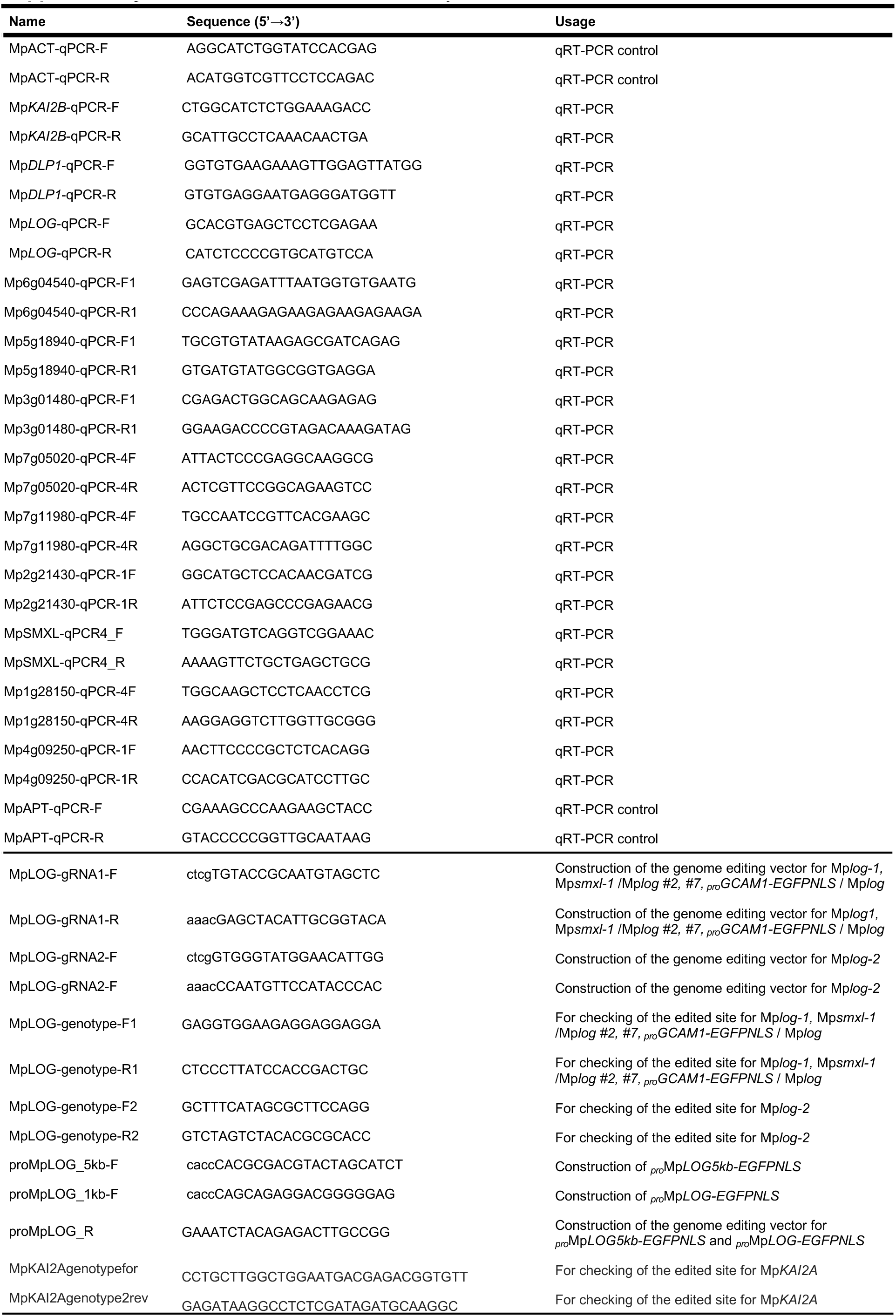
Primers used in this study.

**Supplementary Table 4.** Commonly up-regulated DEGs at 3 or 6 hours treatments.

| id | up | 3h_log2FC | 3h_baseMean | 3h_padj | 6h_log2FC | 6h_baseMean | 6h_padj |
| --- | --- | --- | --- | --- | --- | --- | --- |
| Mp1g00270 | 3h & 6h | 0.71199529 | 1256.64508 | 6.16E-11 | 0.75342512 | 1721.00723 | 3.16E-07 |
| Mp1g05220 | 3h & 6h | 0.32430663 | 457.77033 | 0.04779049 | 0.26343079 | 676.94094 | 0.00219654 |
| Mp1g06800 | 3h & 6h | 0.44440401 | 580.588506 | 0.01521182 | 0.41205929 | 1095.12956 | 2.87E-05 |
| Mp1g09420 | 3h & 6h | 0.55068758 | 65.1325424 | 0.03246633 | 0.47329122 | 109.101509 | 0.01050891 |
| Mp1g12030 | 3h & 6h | 0.20795894 | 1323.42039 | 0.0124496 | 0.21626408 | 2287.16219 | 0.01576324 |
| Mp1g14990 | 3h & 6h | 0.37467467 | 849.265843 | 0.00561349 | 0.29408657 | 1176.74244 | 0.00330979 |
| Mp1g16800 | 3h & 6h | 0.24529377 | 933.39775 | 0.00375365 | 0.38153065 | 1206.00578 | 2.12E-14 |
| Mp1g19760 | 3h & 6h | 0.42194621 | 709.881082 | 1.34E-06 | 0.28379791 | 1011.91202 | 0.00096514 |
| Mp1g20260 | 3h & 6h | 0.57442506 | 614.044222 | 0.00196296 | 0.41296514 | 1263.58154 | 0.00088551 |
| Mp1g20860 | 3h & 6h | 0.55025425 | 107.994477 | 0.00192375 | 0.65203389 | 181.946758 | 7.85E-13 |
| Mp1g23210 | 3h & 6h | 0.43255343 | 1607.15253 | 0.00065748 | 0.52256502 | 1473.46927 | 0.00010732 |
| Mp1g23650 | 3h & 6h | 0.38805047 | 2068.35951 | 0.00022488 | 0.42834882 | 3058.86667 | 3.43E-07 |
| Mp1g26440 | 3h & 6h | 0.29896178 | 2106.2672 | 0.02444225 | 0.27502534 | 2974.70928 | 0.00286682 |
| Mp1g27950 | 3h & 6h | 1.36343192 | 63.3525101 | 7.87E-12 | 1.05034731 | 87.9358105 | 5.99E-15 |
| Mp1g27970 | 3h & 6h | 0.68183383 | 34.0187216 | 0.02049996 | 0.60769779 | 52.6973643 | 0.00314307 |
| Mp1g28150 | 3h & 6h | 0.69169619 | 129.871274 | 0.02656506 | 0.68122185 | 187.161874 | 0.00277994 |
| Mp2g04920 | 3h & 6h | 0.6525162 | 238.402617 | 0.00786532 | 0.66689265 | 452.590131 | 5.61E-12 |
| Mp2g04930 | 3h & 6h | 2.02145468 | 2539.26241 | 5.83E-83 | 2.24479205 | 5023.76135 | 7.37E-98 |
| Mp2g08260 | 3h & 6h | 0.90443304 | 42.2678393 | 8.58E-05 | 1.18137378 | 50.81393 | 4.57E-10 |
| Mp2g11820 | 3h & 6h | 0.36774641 | 267.245838 | 0.04785183 | 0.40397554 | 340.732918 | 0.00918403 |
| Mp2g11860 | 3h & 6h | 0.20642892 | 2943.97609 | 0.03334266 | 0.20173009 | 3929.04504 | 0.00082924 |
| Mp2g12920 | 3h & 6h | 0.38251776 | 3617.3099 | 0.00718084 | 0.20021153 | 5611.11669 | 0.02705655 |
| Mp2g20430 | 3h & 6h | 1.55173848 | 1005.92128 | 0.00076099 | 1.19034491 | 1372.49661 | 5.57E-08 |
| Mp2g21430 | 3h & 6h | 0.86070609 | 754.995099 | 6.97E-10 | 0.80044571 | 1095.74232 | 2.54E-07 |
| Mp2g23680 | 3h & 6h | 0.7005815 | 1358.28972 | 1.06E-11 | 0.49269332 | 2199.47654 | 0.00456028 |
| Mp3g01480 | 3h & 6h | 0.69310861 | 302.597866 | 8.98E-11 | 0.74735789 | 829.597279 | 1.58E-06 |
| Mp3g06310 | 3h & 6h | 0.7513378 | 1835.89488 | 2.90E-21 | 0.71411366 | 2625.65948 | 5.16E-15 |
| Mp3g10080 | 3h & 6h | 0.69077845 | 269.948286 | 3.11E-13 | 0.64467314 | 445.111716 | 9.14E-12 |
| Mp3g12930 | 3h & 6h | 0.65882645 | 463.191219 | 0.00223322 | 0.46602591 | 802.369413 | 1.14E-07 |
| Mp3g14120 | 3h & 6h | 0.70885018 | 245.739077 | 1.13E-08 | 0.68373982 | 320.316212 | 2.30E-08 |
| Mp3g16340 | 3h & 6h | 0.66042476 | 1675.97226 | 0.00555718 | 0.43684369 | 3259.77295 | 0.00808643 |
| Mp3g18590 | 3h & 6h | 0.42627975 | 361.854958 | 0.00409009 | 0.49897105 | 577.383823 | 0.01006989 |
| Mp4g04360 | 3h & 6h | 0.36127223 | 1043.91776 | 1.75E-11 | 0.22069807 | 1708.0943 | 9.26E-05 |
| Mp4g07180 | 3h & 6h | 0.6186193 | 291.765832 | 0.00284636 | 0.47844873 | 368.729709 | 0.0031432 |
| Mp4g09250 | 3h & 6h | 0.73163317 | 341.044842 | 0.00015778 | 1.13246099 | 895.642026 | 4.46E-13 |
| Mp4g10620 | 3h & 6h | 1.22760428 | 63.8126992 | 1.17E-11 | 0.92980761 | 111.798411 | 3.21E-09 |
| Mp4g15090 | 3h & 6h | 0.74310198 | 352.225917 | 9.11E-17 | 0.59018248 | 566.340487 | 1.75E-10 |
| Mp4g17760 | 3h & 6h | 0.34905429 | 474.375869 | 0.00081921 | 0.28902801 | 707.456109 | 0.03357311 |
| Mp4g23700 | 3h & 6h | 0.80511059 | 2407.1577 | 0.00178954 | 0.42409764 | 3428.18882 | 0.01006989 |
| Mp5g02800 | 3h & 6h | 0.91009912 | 352.702567 | 8.48E-05 | 0.7665525 | 470.564821 | 2.58E-12 |
| Mp5g09160 | 3h & 6h | 1.83276982 | 2228.24644 | 5.33E-06 | 2.69106213 | 3907.35782 | 5.45E-64 |
| Mp5g13870 | 3h & 6h | 0.74615607 | 5707.83333 | 0.00231483 | 1.2853932 | 9607.16369 | 8.41E-13 |
| Mp5g18940 | 3h & 6h | 0.65203153 | 271.800019 | 0.03285904 | 1.49107755 | 307.841107 | 1.24E-08 |
| Mp5g21020 | 3h & 6h | 0.37349604 | 452.418094 | 0.03697847 | 0.35520359 | 614.011007 | 0.00669434 |
| Mp5g24030 | 3h & 6h | 0.29005312 | 321.017017 | 0.00629628 | 0.59631715 | 537.816535 | 1.18E-06 |
| Mp6g04540 | 3h & 6h | 1.02928114 | 127.872924 | 1.58E-20 | 1.39009291 | 244.328557 | 3.21E-42 |
| Mp6g05000 | 3h & 6h | 0.51241285 | 5270.17741 | 0.00030862 | 0.48788794 | 9136.97942 | 0.00015082 |
| Mp6g09290 | 3h & 6h | 0.72953826 | 2227.06827 | 6.01E-26 | 0.87299305 | 4632.55725 | 5.03E-20 |
| Mp6g09300 | 3h & 6h | 1.79497269 | 557.666424 | 1.00E-43 | 1.83902871 | 773.837138 | 1.16E-51 |
| Mp6g10520 | 3h & 6h | 1.15679446 | 1325.19115 | 2.31E-06 | 0.67450752 | 2166.74575 | 1.93E-07 |
| Mp6g16970 | 3h & 6h | 0.98569918 | 36.4469729 | 0.00014177 | 0.62057366 | 92.7364342 | 0.01832329 |
| Mp6g17270 | 3h & 6h | 0.46802243 | 718.693599 | 9.86E-05 | 0.33072751 | 1188.18874 | 4.00E-05 |
| Mp7g00770 | 3h & 6h | 0.46670705 | 552.655686 | 0.00024196 | 0.44415537 | 527.834293 | 0.00430972 |
| Mp7g02090 | 3h & 6h | 0.26584056 | 445.00001 | 0.00115205 | 0.36544247 | 796.678807 | 8.90E-09 |
| Mp7g03280 | 3h & 6h | 0.15017403 | 6658.39585 | 0.00106976 | 0.1557989 | 9666.96612 | 0.00078739 |
| Mp7g05020 | 3h & 6h | 1.15624989 | 224.312854 | 5.09E-08 | 0.52585241 | 396.322755 | 0.0107156 |
| Mp7g05040 | 3h & 6h | 0.38193268 | 1002.42753 | 0.00010909 | 0.26556569 | 1351.89649 | 0.00250252 |
| Mp7g06170 | 3h & 6h | 0.41777761 | 1539.04423 | 0.03460412 | 0.32471072 | 2778.47281 | 0.01775762 |
| Mp7g06960 | 3h & 6h | 0.24023941 | 611.2978 | 0.01946749 | 0.25942147 | 965.651725 | 1.79E-06 |
| Mp7g10170 | 3h & 6h | 0.71906914 | 433.444427 | 0.00029243 | 0.33534545 | 601.181451 | 0.00411053 |
| Mp7g11980 | 3h & 6h | 0.59849091 | 244.782877 | 0.00036222 | 0.65215072 | 519.034127 | 0.00055537 |
| Mp7g12790 | 3h & 6h | 0.47675327 | 643.597742 | 4.49E-05 | 0.56346513 | 1500.6234 | 1.19E-09 |
| Mp7g15820 | 3h & 6h | 0.45370888 | 211.36092 | 0.00099343 | 0.31242873 | 233.252618 | 0.03668743 |
| Mp7g17880 | 3h & 6h | 0.63342812 | 322.633469 | 0.00136036 | 0.5163001 | 458.748319 | 0.03219593 |
| Mp7g19700 | 3h & 6h | 0.29593891 | 300.868389 | 0.02560661 | 0.29757352 | 632.310843 | 0.00038149 |
| Mp8g02280 | 3h & 6h | 0.64121613 | 138.831002 | 0.00039438 | 1.03876754 | 134.832233 | 5.68E-13 |
| Mp8g02500 | 3h & 6h | 0.6556798 | 61.5528148 | 3.66E-05 | 0.37690322 | 115.295472 | 0.0259067 |
| Mp8g04780 | 3h & 6h | 0.46581611 | 2993.91415 | 5.87E-05 | 0.46257829 | 3181.2575 | 9.68E-23 |
| Mp8g04910 | 3h & 6h | 0.31366805 | 523.976168 | 2.80E-05 | 0.48308021 | 895.365066 | 4.55E-08 |
| Mp8g04950 | 3h & 6h | 0.30984732 | 706.877142 | 0.00254352 | 0.25439954 | 1235.69319 | 0.00063011 |
| Mp8g06260 | 3h & 6h | 0.56969138 | 163.541687 | 0.00784358 | 0.78941303 | 292.470683 | 2.48E-13 |
| Mp8g06760 | 3h & 6h | 0.57411042 | 230.303056 | 0.00118944 | 0.34436101 | 338.2014 | 0.04759426 |
| Mp8g08780 | 3h & 6h | 0.45675556 | 356.245186 | 0.00480768 | 0.42412736 | 482.935267 | 7.83E-10 |
| Mp8g10420 | 3h & 6h | 0.24221534 | 369.267014 | 0.0143674 | 0.25348626 | 532.024784 | 0.00087073 |
| Mp8g14310 | 3h & 6h | 0.18747246 | 668.221672 | 0.02535938 | 0.20937797 | 1224.15478 | 0.04056111 |
| Mp8g19020 | 3h & 6h | 0.56776246 | 488.970528 | 8.13E-07 | 0.60704447 | 552.199446 | 1.53E-08 |

## References

1. Steeves T. A. &, Sussex I. M. Patterns in Plant Development. (1989).

2. Frey W. & Kürschner H. Asexual reproduction, habitat colonization and habitat maintenance in bryophytes. Flora - Morphology, Distribution, Functional Ecology of Plants 206, 173–184 (2011).

3. Ishida S. et al. Diminished auxin signaling triggers cellular reprogramming by inducing a regeneration factor in the liverwort *Marchantia polymorpha*. Plant Cell Physiol 63, 384–400 (2022).

4. Ikeuchi M. et al. Molecular mechanisms of plant regeneration. Annu Rev Plant Biol 70, 377–406 (2019).

5. Shimamura M. Marchantia polymorpha: taxonomy, phylogeny and morphology of a model system. Plant Cell Physiol 57, 230–256 (2016).

6. Burgeff H. Genetische studien an Marchantia. Einführung einer neuen Pflanzenfamilie in die genetische Wissenschaft. Ficher, (1943).

7. Kato H., Yasui Y. & Ishizaki K. Gemma cup and gemma development in Marchantia polymorpha. New Phytol 228, 459–465 (2020).

8. Waadt R. et al. Plant hormone regulation of abiotic stress responses. Nat Rev Mol Cell Biol 23, 680–694 (2022).

9. Gray W. M. Hormonal regulation of plant growth and development. PLoS Biol 2, E311 (2004).

10. Vanstraelen M. & Benkova E. Hormonal interactions in the regulation of plant development. Annu Rev Cell Dev Biol 28, 463–487 (2012).

11. Komatsu A. et al. Control of vegetative reproduction in Marchantia polymorpha by the KAI2-ligand signaling pathway. Curr Biol 33, 1196–1210 e1194 (2023).

12. Mizuno Y. et al. Major components of the KARRIKIN INSENSITIVE2-dependent signaling pathway are conserved in the liverwort Marchantia polymorpha. Plant Cell 33, 2395–2411 (2021).

13. Nelson D. C. et al. F-box protein MAX2 has dual roles in karrikin and strigolactone signaling in Arabidopsis thaliana. Proc Natl Acad Sci USA 108, 8897–8902 (2011).

14. Waters M. T. et al. Specialisation within the DWARF14 protein family confers distinct responses to karrikins and strigolactones in Arabidopsis. Development 139, 1285–1295 (2012).

15. Conn C. E. & Nelson D. C. Evidence that KARRIKIN-INSENSITIVE2 (KAI2) receptors may perceive an unknown signal that is not karrikin or strigolactone. Front Plant Sci 6, 1219 (2015).

16. Soundappan I. et al. SMAX1-LIKE/D53 family members enable distinct MAX2-dependent responses to strigolactones and karrikins in Arabidopsis. Plant Cell 27, 3143–3159 (2015).

17. Waters M. T. & Nelson D. C. Karrikin perception and signalling. New Phytol 237, 1525–1541 (2023).

18. Zheng J. et al. Karrikin signaling acts parallel to and additively with strigolactone signaling to regulate rice mesocotyl elongation in darkness. Plant Cell 32, 2780–2805 (2020).

19. Wang L. et al. Strigolactone and karrikin signaling pathways elicit ubiquitination and proteolysis of SMXL2 to regulate hypocotyl elongation in Arabidopsis. Plant Cell 32, 2251–2270 (2020).

20. Stanga J. P. et al. SUPPRESSOR OF MORE AXILLARY GROWTH2 1 controls seed germination and seedling development in Arabidopsis. Plant Physiol 163, 318–330 (2013).

21. Stanga J.P., Morffy N. & Nelson D. C. Functional redundancy in the control of seedling growth by the karrikin signaling pathway. Planta 243, 1397–1406 (2016).

22. Wang L. et al. Strigolactone signaling in Arabidopsis regulates shoot development by targeting D53-Like SMXL repressor proteins for ubiquitination and degradation. Plant Cell 27, 3128–3142 (2015).

23. Wang L. et al. Transcriptional regulation of strigolactone signalling in Arabidopsis. Nature 583, 277–281 (2020).

24. Song X. et al. IPA1 functions as a downstream transcription factor repressed by D53 in strigolactone signaling in rice. Cell Res 27, 1128–1141 (2017).

25. Hu J. et al. BES1 functions as the co-regulator of D53-like SMXLs to inhibit *BRC1* expression in strigolactone-regulated shoot branching in *Arabidopsis*. Plant Commun 1, 100014 (2019).

26. Xie, Y. et al. *Arabidopsis* FHY3 and FAR1 integrate light and strigolactone signaling to regulate branching. Nat Commun 11, 1955. (2020).

27. Wang L., Waters M. T. & Smith S. M. Karrikin-KAI2 signalling provides Arabidopsis seeds with tolerance to abiotic stress and inhibits germination under conditions unfavourable to seedling establishment. New Phytol 219, 605–618 (2018).

28. Xu P., Jinbo H. & Cai W. Karrikin signaling regulates hypocotyl shade avoidance response by modulating auxin homeostasis in Arabidopsis. New Phytol 236, 1748–1761 (2022).

29. Li W. et al. The karrikin receptor KAI2 promotes drought resistance in Arabidopsis thaliana. PLoS Genet 13, e1007076 (2017).

30. Nelson D. C. et al. Karrikins enhance light responses during germination and seedling development in Arabidopsis thaliana. Proc Natl Acad Sci USA 107, 7095–7100 (2010).

31. Choi J. et al. The negative regulator SMAX1 controls mycorrhizal symbiosis and strigolactone biosynthesis in rice. Nat Commun 11, 2114 (2020).

32. Varshney K. & Gutjahr C. KAI2 can do: karrikin receptor function in plant development and response to abiotic and Biotic Factors. Plant Cell Physiol 64, 984–995 (2023).

33. Villaecija-Aguilar J. A. et al. KAI2 promotes Arabidopsis root hair elongation at low external phosphate by controlling local accumulation of AUX1 and PIN2. Curr Biol 32, 228–236 e223 (2022).

34. Carbonnel S. et al. The karrikin signaling regulator SMAX1 controls Lotus japonicus root and root hair development by suppressing ethylene biosynthesis. Proc Natl Acad Sci USA 117, 21757–21765 (2020).

35. Hamon-Josse M. et al. KAI2 regulates seedling development by mediating light-induced remodelling of auxin transport. New Phytol 235, 126–140 (2022).

36. Brun G. et al. CYP707As are effectors of karrikin and strigolactone signalling pathways in Arabidopsis thaliana and parasitic plants. Plant Cell Environ 42, 2612–2626 (2019).

37. Bowman J. L. et al. Insights into land plant evolution garnered from the Marchantia polymorpha genome. Cell 171, 287–304 e215 (2017).

38. Bonhomme S. & Guillory A. Synthesis and signalling of strigolactone and KAI2-ligand signals in bryophytes. J Exp Bot 73, 4487–4495 (2022).

39. Kodama K. et al. An ancestral function of strigolactones as symbiotic rhizosphere signals. Nat Commun 13, 3974 (2022).

40. Lopez-Obando M. et al. Physcomitrella patens MAX2 characterization suggests an ancient role for this F-box protein in photomorphogenesis rather than strigolactone signalling. New Phytol 219, 743–756 (2018).

41. Bythell-Douglas R. et al. Evolution of strigolactone receptors by gradual neo-functionalization of KAI2 paralogues. BMC Biol 15, 52 (2017).

42. Kodama K., Xie, X. & Kyozuka J. The D14 and KAI2 orthologs of gymnosperms sense strigolactones and KL mimics, respectively, and the signals are transduced to control downstream genes. Plant Cell Physiol 64, 1057–1065 (2023)

43. Conn C. E., et al. PLANT EVOLUTION. Convergent evolution of strigolactone perception enabled host detection in parasitic plants. Science 349, 540–543 (2015).

44. Toh S. et al. Structure-function analysis identifies highly sensitive strigolactone receptors in Striga. Science 350, 203–207 (2015).

45. Lopez-Obando M. et al. The Physcomitrium (Physcomitrella) patens PpKAI2L receptors for strigolactones and related compounds function via MAX2-dependent and -independent pathways. Plant Cell 33, 3487–3512 (2021).

46. Kurakawa T. et al. Direct control of shoot meristem activity by a cytokinin-activating enzyme. Nature 445, 652–655 (2007).

47. Yasui Y. et al. GEMMA CUP-ASSOCIATED MYB1, an ortholog of axillary meristem regulators, is essential in vegetative reproduction in Marchantia polymorpha. Curr Biol 29, 3987–3995.e3985 (2019).

48. Hirose N. et al. Regulation of cytokinin biosynthesis, compartmentalization and translocation. J Exp Bot 59, 75–83 (2008).

49. Kamada-Nobusada T. & Sakakibara H. Molecular basis for cytokinin biosynthesis. Phytochemistry 70, 444–449 (2009).

50. Aki S. S. et al. Cytokinin signaling is essential for organ formation in *Marchantia polymorpha*. Plant Cell Physiol 60, 1842–1854 (2019).

51. Muller D., Schmitz G. & Theres K. Blind homologous R2R3 Myb genes control the pattern of lateral meristem initiation in Arabidopsis. Plant Cell 18, 586–597 (2006).

52. Aki S.S. et al. R2R3-MYB transcription factor GEMMA CUP-ASSOCIATED MYB1 mediates the cytokinin signal to achieve proper organ development in Marchantia polymorpha. Sci Rep 12, 21123 (2022).

53. Hong L. & Fletcher J. C. Stem cells: engines of plant growth and development. Int J Mol Sci 24, (2023).

54. Chickarmane V. S. et al. Cytokinin signaling as a positional cue for patterning the apical-basal axis of the growing Arabidopsis shoot meristem. Proc Natl Acad Sci USA 109, 4002–4007 (2012).

55. Wang X. et al. Evolution and roles of cytokinin genes in angiosperms 1: Do ancient IPTs play housekeeping while non-ancient IPTs play regulatory roles? Hortic Res 7, 28 (2020).

56. Nishii K. et al. Tangled history of a multigene family: The evolution of ISOPENTENYLTRANSFERASE genes. PLoS One 13, e0201198 (2018).

57. Lindner A. C. et al. Isopentenyltransferase-1 (IPT1) knockout in Physcomitrella together with phylogenetic analyses of IPTs provide insights into evolution of plant cytokinin biosynthesis. J Exp Bot 65, 2533–2543 (2014).

58. Naseem M., Sarukhanyan E. & Dandekar T. LONELY-GUY knocks every door: crosskingdom microbial pathogenesis. Trends Plant Sci 20, 781–783 (2015).

59. Samanovic M. I. et al. Proteasomal control of cytokinin synthesis protects Mycobacterium tuberculosis against nitric oxide. Mol Cell 57, 984–994 (2015).

60. Busch B. L. et al. Shoot branching and leaf dissection in tomato are regulated by homologous gene modules. Plant Cell 23, 3595–3609 (2011).

61. Keller T. et al. Arabidopsis REGULATOR OF AXILLARY MERISTEMS1 controls a leaf axil stem cell niche and modulates vegetative development. Plant Cell 18, 598–611 (2006).

62. Schmitz G. et al. The tomato Blind gene encodes a MYB transcription factor that controls the formation of lateral meristems. Proc Natl Acad Sci USA 99, 1064–1069 (2002).

63. Umehara M. et al. Inhibition of shoot branching by new terpenoid plant hormones. Nature 455, 195–200 (2008).

64. Gomez-Roldan V. et al. Strigolactone inhibition of shoot branching. Nature 455, 189–194 (2008).

65. Dun E. A. et al. Strigolactones and shoot branching: what is the real hormone and how does it work? Plant Cell Physiol 64, 967–983 (2023).

66. Waters M. T., et al. Strigolactone signaling and evolution. Annu Rev Plant Biol 68, 291–322 (2017).

67. Walker C.H. et al. Strigolactone synthesis is ancestral in land plants, but canonical strigolactone signalling is a flowering plant innovation. BMC Biol 17, 70 (2019).

68. von Schwartzenberg K. Moss biology and phytohormones--cytokinins in *Physcomitrella*. Plant biol 8, 382–388 (2006).

69. Guillory A. & Bonhomme S. Phytohormone biosynthesis and signaling pathways of mosses. Plant Mol Biol 107, 245–277 (2021).

70. Li L. et al. Molecular regulation and evolution of cytokinin signaling in plant abiotic stresses. Plant Cell Physiol 63, 1787–1805 (2022).

71. Tokunaga H. et al. Arabidopsis lonely guy (LOG) multiple mutants reveal a central role of the LOG-dependent pathway in cytokinin activation. Plant J 69, 355–365 (2012).

72. Kojima M. et al. A cell wall-localized cytokinin/purine riboside nucleosidase is involved in apoplastic cytokinin metabolism in Oryza sativa. Proc Natl Acad Sci USA 120, e2217708120 (2023).

73. Love M. I., Huber W. & Anders S. Moderated estimation of fold change and dispersion for RNA-seq data with DESeq2. Genome Biol 15, 550 (2014).

74. Yu G. et al. clusterProfiler: an R package for comparing biological themes among gene clusters. OMICS 16, 284–287 (2012).

75. Sugano S. S. et al. Efficient CRISPR/Cas9-based genome editing and its application to conditional genetic analysis in Marchantia polymorpha. PLoS One 13, e0205117 (2018).

76. Kojima M. et al. Highly sensitive and high-throughput analysis of plant hormones using MS-probe modification and liquid chromatography-tandem mass spectrometry: an application for hormone profiling in Oryza sativa. Plant Cell Physiol 50, 1201–1214 (2009).

